# Drivers of adaptive capacity in wild populations: implications for genetic interventions

**DOI:** 10.1101/2021.02.25.432972

**Authors:** G Torda, K Quigley

## Abstract

The unprecedented rate of environmental change in the Anthropocene poses evolutionary challenges for wild populations globally. Active human interventions are being increasingly considered to accelerate natural adaptive processes. Evolutionary models can evaluate how species may fare under future climate, elucidate which evolutionary processes are critical to rapid adaptation, and how active interventions may influence fitness trajectories of organisms. Here we use polygenic metapopulation adaptation models to quantify the relative importance (effect sizes) of different eco-evolutionary parameters on the rates of adaptation in wild populations i) without active interventions, and ii) under a subset of active interventions. We demonstrate that genetic diversity (heterozygosity, He), population connectivity and the effect size of additive genetic variance are the primary drivers of natural adaptation rates. We quantify the effect sizes of these parameters on population fitness across three proposed assisted evolution scenarios and identify critical thresholds for intervention effectiveness and implementation. Specifically, the interventions tested here were most effective at low levels of genetic diversity in target populations (He < 0.2) and when timed during a cold-to-warm phase of an ENSO-like oscillation. Beneficial levels of connectivity were highly dependent on desired outcomes for the meta-population. We also present a global meta-analysis of genetic diversity in tropical reef-building corals as a case study of how thresholds derived from evolutionary models can be used to guide decision making by managers. We find genetic diversity to be highly variable by coral taxon and region, highlighting how thresholds from evolutionary models can be used in conjunction with empirical data to assess intervention needs and priorities. Finally, we highlight the critical knowledge and data gaps to produce the next suite of applied models for conservation management decision-support.

## Main

Ecosystems worldwide are degrading at an alarming rate as species struggle to keep pace with novel environmental pressures arising from rapid climate change. There is growing concern that keystone species that maintain ecosystem functioning will be unable to match the speed of temperature change by natural adaptive processes. The loss of such keystone species may lead to further ecosystem collapse and a rapid transition into degraded alternate stable states ^1^ that fail to provide ecosystem services. Assisted evolution methods aim to preserve or restore ecosystems by enhancing adaptive traits of species most critically threatened by climate change ^2–4^. To assess the feasibility of success and the risks of these genetic interventions in the wild, the main factors driving adaptive capacities need to be assessed and quantified.

Traits can be determined by one to multiple genes, and these causal variants can generally be described as quantitative trait loci (QTLs). The genetic basis of polygenic trait evolution is complex ^5^. Key parameters like the number of genes and magnitude of the effect of each gene (effect size) are still unknown for a majority of QTLs that code fitness-related traits^6^, as well as the influence of the interaction of genes and traits that drive adaptation^7^. Predictive evolutionary simulations (e.g. ^8^) can be used to explore spatial and temporal scales of adaptation and model the relative importance of a wide range of ecological and genetic population parameters that may drive adaptive capacities. Combined with empirical data, this allows for a powerful approach to assess the adaptive potential of natural populations and better predict the potential outcomes of different intervention methods. Quantifying and incorporating these evolutionary dynamics into managing for climate change is essential for both passive and active conservation planning ^9^.

The evolution of traits within populations may require many generations, although examples exist that demonstrate rapid evolution is possible under particular scenarios, reviewed in ^10,11^. Rapid adaptation to ecological change occurs through the exploitation of existing “standing” genetic variation within populations. Indeed, the importance of genetic diversity, and its maintenance for protecting populations from extinction, or alternatively, rescuing them from population decline, has long been recognized in terrestrial and marine environments (e.g. the Florida panther ^12^, the arctic fox ^13^, and the mountain pygmy possum ^14^). The manipulation of genetic diversity is therefore an important target for practitioners undertaking the assisted evolution of captive and wild populations, but the scope of impact as well as the benefits and risks of different interventions are still unknown, especially in species with large population sizes and variable genetic diversity ^9^. Here we present a series of evolutionary simulations that encompass a range from single-gene-single-population models of adaptation to multiple-genes-multiple-populations models (hereafter polygenic metapopulation models) and provide an assessment of the effect of multiple ecological and genetic parameters (including genetic diversity) on the rate and potential scope of adaptation (i.e. the range of different adaptive capacities). We then discuss the relative importance of these parameters in determining the effects of three of the currently >40 recognised interventions (both genetic and non-genetic) ^15,16^ proposed on one ecosystem, coral reefs. Broadly, they relate to the enhancement of particular traits of the host (e.g. heat tolerance) via genetic selection, one of four broad intervention categories being considered ^2^. Finally, we present a global meta-analysis of empirically derived data on the genetic diversity of tropical reef-building corals. We use this data to demonstrate the application of the thresholds of genetic diversity derived from evolutionary models to aid in decision-making for genetic interventions.

## Results

### The relative importance of model parameters in shaping adaptive capacities

The model parameters and assumptions explored are described in detail in the Methods. Briefly, connectivity among populations followed a classic stepping-stone meta-popuation model (Fig. 4), in which a temperature gradient was applied such that the environment of population 1 was warmer (1.1°C) than the environment of population 5 (0.9°C). Temperature was held stable for 100 generations (a “burn-in” period), after which an El Niño Southern Oscillation (ENSO) – like cycle with an amplitude of 0.1°C and a warming period (wavelength) of five generations was slowly incorporated over 100 generations. Warming was simulated by incorporating a temperature increase of 0.01°C per generation, starting at generation 200, and lasting 150 generations (further information in Methods). Environmental tolerance was modelled as the width of the fitness function (Fig. 5). To quantify the importance of key model parameters (genetic diversity, number of genes coding the trait, environmental tolerance, connectivity, mutation rate, population size) for determining adaptive capacities of populations, we calculated two metrics: (i) the rate of adaptation as defined by fitness change over the first 20 years of warming (zero indicating no fitness change in a warming environment, hence rapid adaptation); and (ii) mean population fitness achieved after 20, 50 and 100 generations of warming.

#### i) Factors influencing the rate of adaptation

The most influential parameter for determining the rate of adaptation in wild populations was the number of genes underpinning the trait (Fig. 1A; standard deviation of fitness difference between the start of warming and 20 generations later, across ten independent simulations 0.22 ± 0.02). Adaptation was slower for traits encoded by a higher number of genes;the rate of adaptation almost perfectly matched the rate of evironmental change for traits encoded by 50 loci (−0.008 ± 0.006 fitness change over 20 years), while 1,000-loci-traits suffered a significant drop in fitness over the same period (−0.51 ± 0.03 fitness change; Supplementary Fig. 8).

**Figure 1.**
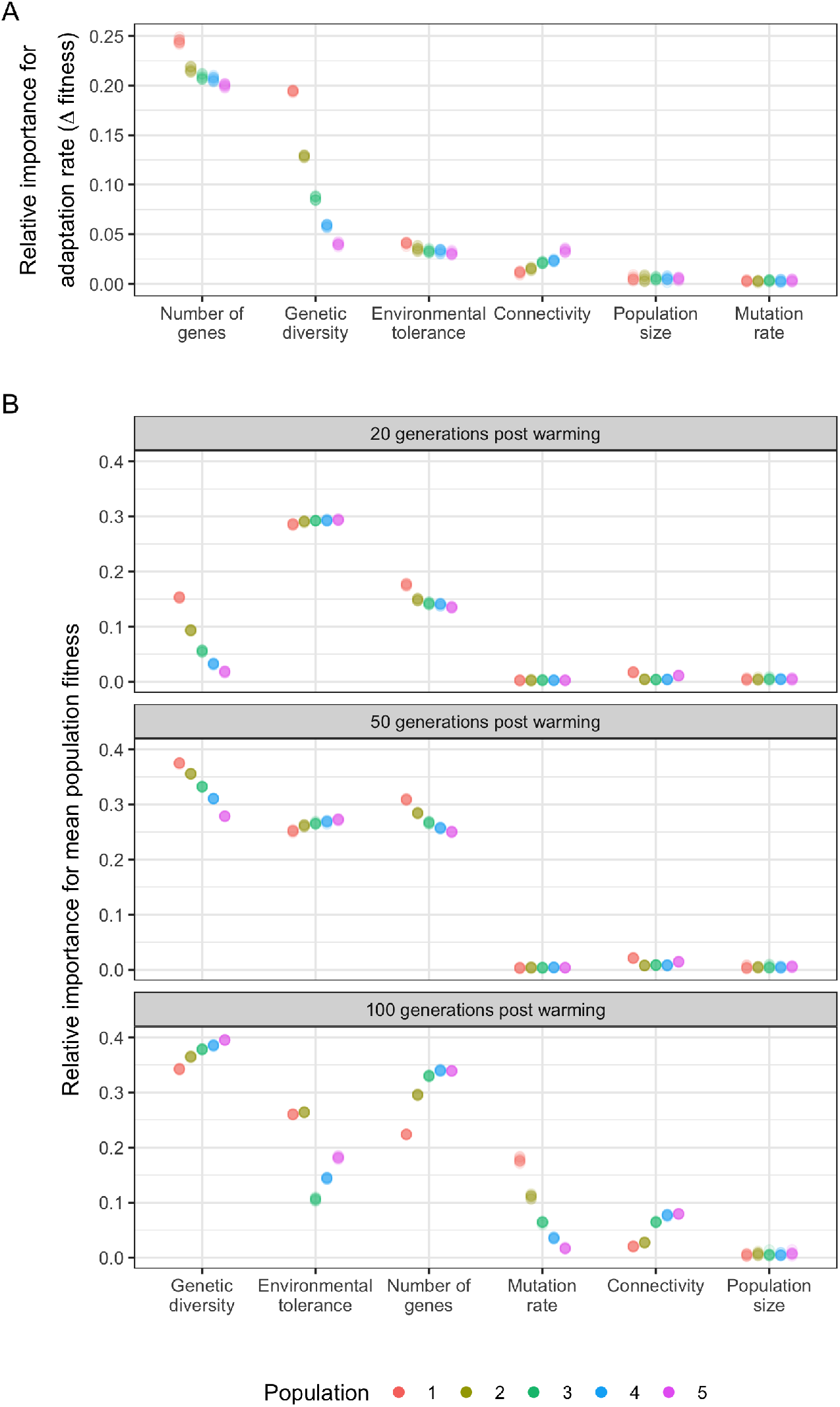
The relative importance of population genomic parameters for the (A) rate of adaptation (change in fitness in the first 20 generations of warming; and (B) mean population fitness after 20, 50 and 100 generations of warming. Relative importance of model parameters for the rate of adaptation was measured by the standard deviation (sd) of the fitness differences at the start of a 0.01°C/generation warming and 20 generations later, varying the target parameter while keeping the others constant. Relative importance of model parameters on mean population fitness was measured by the sd of the fitness values at 20, 50 and 100 generations of 0.01°C/generation warming, varying the target parameter while keeping the others constant. Populations were distributed along an environmental gradient with decreasing temperature from Population 1 to 5, and with twice as much migration from warm to cold populations than from cold to warm populations. Results for ten independent simulations are presented for each permutation of parameters as raw data (due to low within-group variability) and overlayed as points with transparency.

The second most influential parameter for determining the rate of adaptation was genetic diversity, measured by heterozygosity (standard deviation of fitness difference among simulations 0.10 ± 0.05), especially in the warmest population (i.e. closest to the thermal maximum of the metapopulation; 0.18 ± 0.001.; Fig. 1A). Adaptation rate increased with genetic diversity in all populations following a steep saturation curve (Supplementary Fig. 8). Below the threshold value of He=0.06, adaptation was severley compromised.

Metapopulation connectivity (0.02 ± 0.007) and the width of the environmental tolerance curve (0.03 ± 0.004) also influenced rates of adaptation as measured by fitness change in the first 20 years of warming. Connectivity influenced the rate of adaptation most in the coldest population (i.e population downstream from all other populations; 0.03 ± 0.001; Fig. 1A). Increasing connectivity resulted in higher rates of adaptation in all populations but the warmest, where the influx of maladpative gene variants for that population resulted in slower adaptation. Beyond a critical value of connectivity (10% for population 5, 20% for populations 3 and 4), the population fitness increased compared to pre-warming levels (Supplementary Fig. 8). Higher environmental tolerance generally resulted in slower rates of adaptation until a threshold value was reached beyond which further increases in environmental tolerance did not result in a decrease of adaptation rates (Supplementary Fig. 8). This threshold value was 10% of the maximum theoretical phenotypic trait value, i.e. 0.2 in our main simulations (see Supplementary material for re-scaling the parameter values).

#### ii) Factors influencing changes to mean population fitness

The parameters with the strongest influence on mean population fitness after 20 generations of warming were environmental tolerance (standard deviation of fitness values across simulations 0.29 ± 0.003), the number of genes (0.15 ± 0.01), and genetic diversity (0.07 ± 0.05; Fig. 1B). In general, high environmental tolerance (i.e. an increased width of the fitness function) resulted in overall higher fitness, but also allowed for higher genotype-environment mismatch. The higher number of genes encoding a trait, the lower the mean population fitness was after 20 generations of warming; while greater genetic diversity resulted in higher fitness, reaching an asymptote (saturation) at heterozygosity values between 0.06 and 0.20 (Supplementary Fig. 9).

Adaptive capacity came close to exhausted after 50 generations of warming (Fig. 4). As warming progressed, the relative importance of genetic diversity and the number of genes influencing fitness increased for all populations (0.33 ±0.03 and 0.27 ± 0.02, respectively); while environmental tolerance remained highly influential (0.26 ± 0.01; Fig. 1B). After 100 generations of warming, and as temperatures approached the theoretically possible thermal maximum in the simulations (range 1.2-2.4°C, depending on mutation effect size and the number of loci; Fig. 4), genetic diversity again was the most influential parameter determining fitness across all five populations given the modelled genetic architectures. The second most influential parameter was the number of genes (Fig. 1B, Supplementary Fig. 9). For the warmest two populations, environmental tolerance was also highly influential after 100 generations of warming (0.26 ± 0.001 for both populations 1 and 2). Although novel mutations were rare (maximum mutation rate of 10e^-^5 over 100 QTLs), after 100 generations new mutations increased fitness as populations approached their limits of adaptation based on standing genetic variation alone (Supplementary Fig. 9). Mutation rate therefore became an influential parameter in the warmest population (0.18 ± 0.004), with decreasing effect in cooler populations (Fig. 1B) after 100 generations of warming.

### The relative importance of model parameters for the outcome of genetic interventions

Three genetic intervention scenarios where simulated to model their effect on the fitness trajectory of wild populations, as described in detail in the Methods. Briefly, scenario 1 simulated the use of “interpopulation hybrids” produced by reproductively crossing individuals from populations 1 and 5 for inoculation to population 4 (a cooler than average population); scenario 2 redistributed the standing genetic variation of the metapopulation to create inocula (i.e. offspring added to the receiving population) with phenotypes that maximized fitness for the environment of population 4; and finally, scenario 3 increased the standing genetic variation with the addition of new alleles to create inocula with optimal phenotypes that maximized fitness to the environment of the receiving population.

Across the three types of simulated interventions, the most influential biological parameters were the number of genes coding the trait (standard deviation of fitness effect of simulations with different parameter values 0.12 ± 0.06) and genetic diversity (0.06 ± 0.02), followed by environmental tolerance (0.04 ± 0.006) and connectivity (0.03 ± 0.003; Fig. 2B). Interventions that did not create novel diversity, i.e. those that created inocula harnessing only the standing genetic variation of the metapopulation (scenarios 1 and 2) resulted in only temporary fitness effects of generally < 10 generations. Alternatively, inocula that carried novel genetic variation to the metapopulation (scenario 3) created long lasting fitness effects (Fig. 3; Supplementary Figs. 10-14).

**Figure 2.**
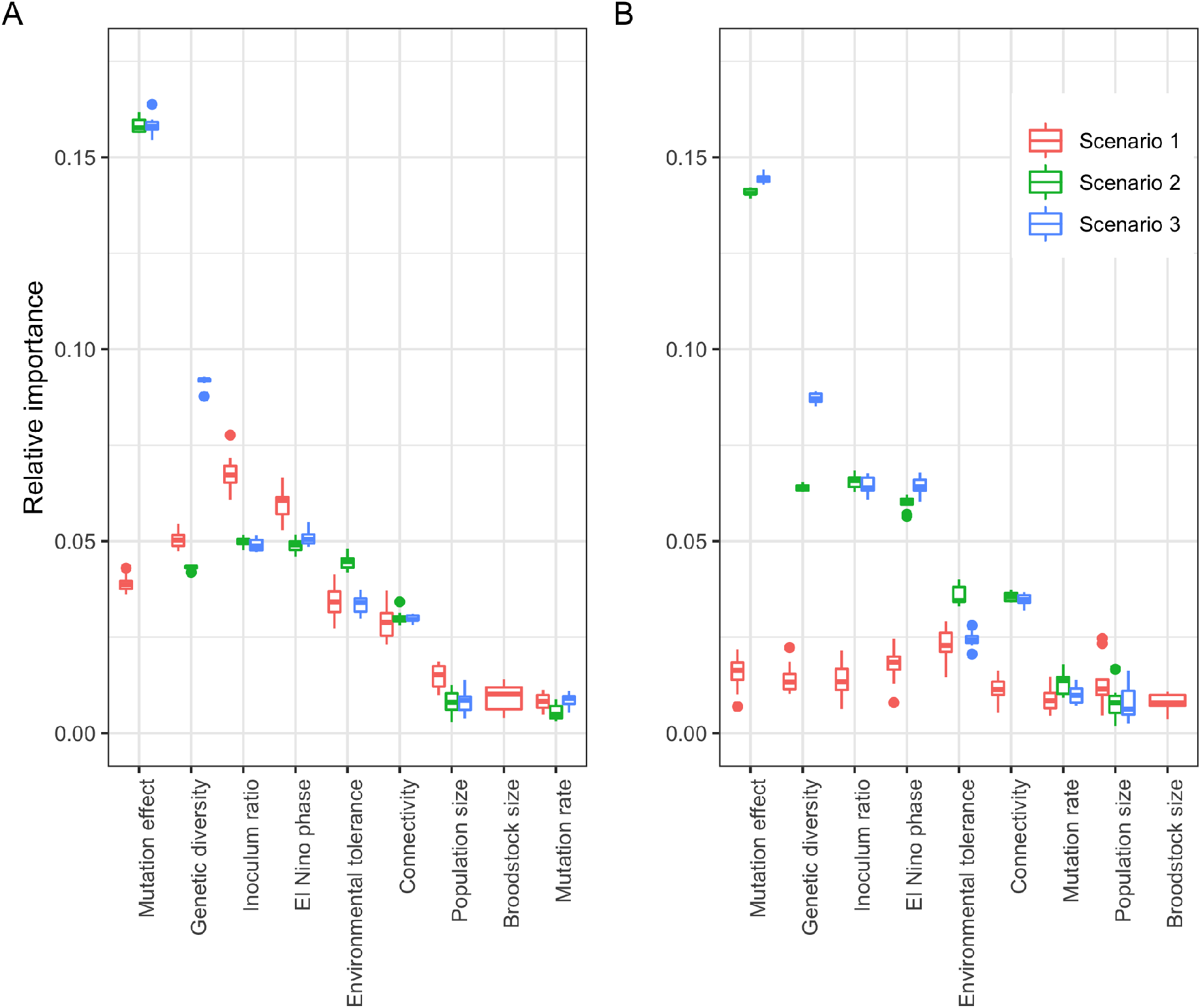
Relative importance of model parameters on the fitness effect of three intervention types: reproductively crossing extreme warm and cold populations (scenario 1), reshuffling the standing genetic variation artificially (scenario 2), or adding novel genetic variation (scenario 3). Relative importance was measured by the standard deviation of fitness effect, varying the target parameter while keeping the others constant. When the target population lies in the middle of the environmental gradient that exists along the range of the metapopulation (Panel B), Scenario 1 (crossing warm and cold populations) has negligible fitness effect; while the same intervention in a colder population yields fitness effect (Panel A). Boxplots show the results of ten independent simulations for each parameter combination.

**Figure 3.**
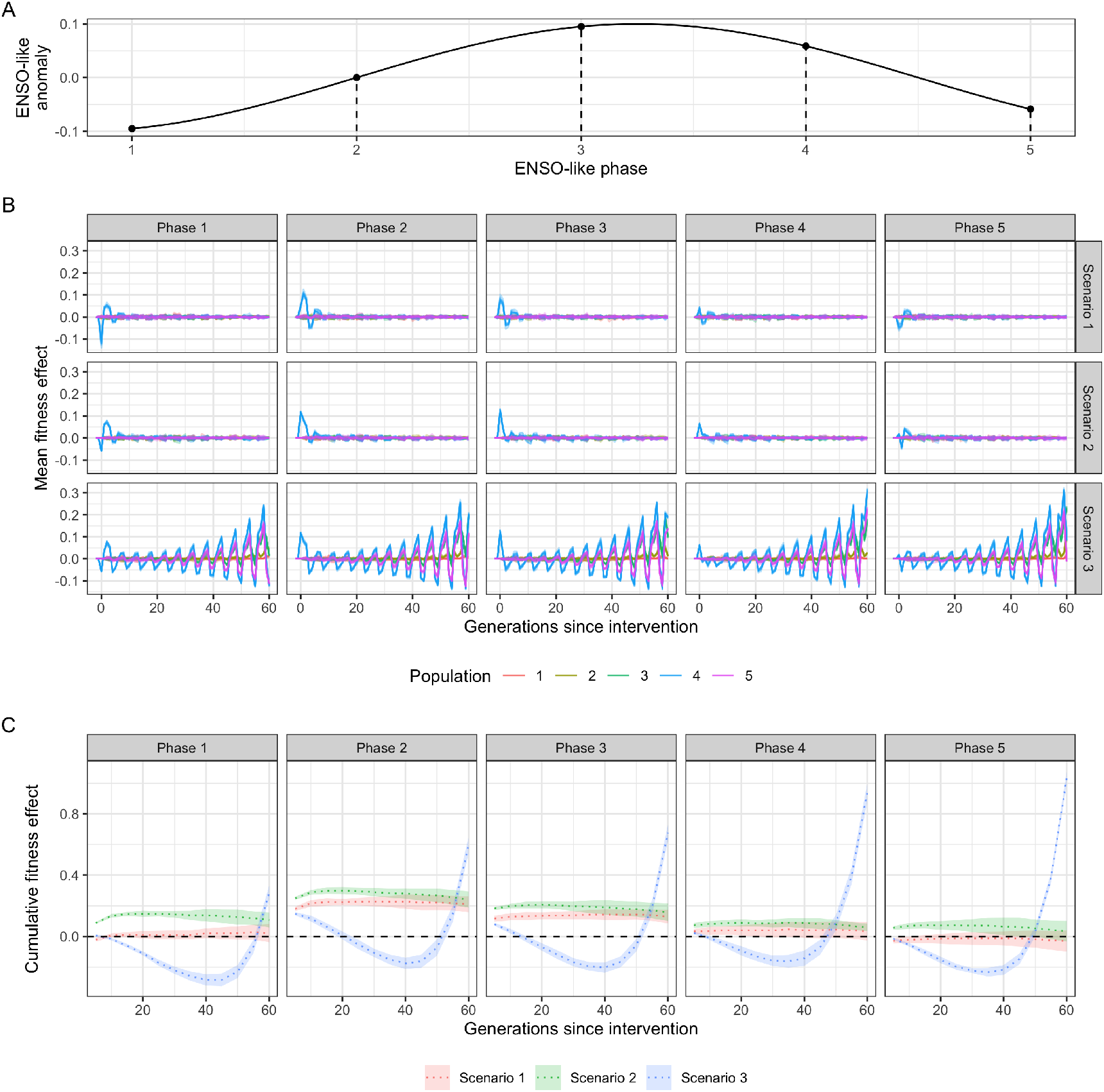
(A) Schematic of the ENSO-like anomalies; (B) the influence of the timing of interventions in relation to ENSO-like phases on fitness effect; and (C) the cumulative fitness effect (over generations) of interventions in the target population (population 4) at different ENSO-like phases. Scenario 1 = crossing warm and cold populations; scenario 2 = re-shuffling the standing genetic variation within the metapopulation; scenario 3 = introducing novel genetic variation to the metapopulation.

#### Number of genes

The effect of interventions on fitness was larger when the trait was encoded by a higher number of genes (Supplementary Fig. 12), because the rate of natural adaptation is lower for highly polygenic traits (see above; Supplementary Fig. 8A), and therefore populations with highly polygenic traits were chronically maladapted at the time of interventions. For example, the fitness effect in the second generation after intervention was 51-fold higher when the trait was encoded by 1,000 loci compared to 50 loci (scenario 2).

#### Genetic diversity

Under extreme low levels of genetic diversity (He < 0.06), the introduction of novel genetic variation (scenario 3) resulted in a strong, positive fitness effect. In contrast, interventions that utilized only the existing standing genetic variation of the metapopulation (scenarios 1 and 2) had negligible fitness effects on the long-term (Supplementary Fig. 10). As genetic diversity increased in the metapopulation, the immediate fitness effect of adding novel genetic variation decreased, while intervention scenarios 1 and 2 created a transient “fitness ripple” (Supplementary Fig. 10). For example, in a low genetic diversity metapopulation (He = 0.06), the immediate fitness effect of intervention scenario 3 was nine-times higher than in a high genetic diversity metapopulation (He = 0.48); and the mean fitness effect in the first ten generations was 0.24 units higher (Supplementary Fig. 10).

#### Connectivity

High rates of connectivity decreased the magnitude of each intervention scenario on the fitness effects in the target population. For example, reproductively crossing extreme populations to obtain inocula (scenario 1) resulted in an immediate fitness benefit of 0.08 units in target population 4 at a 0.01% connectivity rate; which was 6-times higher than the 0.014 units fitness benefit at a 33% connectivity rate (Supplementary Fig. 11). Introduced novel genetic variation (scenario 3) spread faster among populations when connectivity was high. The fitness effect of intervention scenario 3 on population 5 (downstream from the intervened population 4) was highest at an intermediate (1%) connectivity rate. For example, two generations after the intervention, the fitness benefit was 9-times higher at 1% connectivity rate than at 0.1%; and 24-times higher at 1% connectivity rate than at 10% in population 5. For upstream populations, fitness benefit became significant only tens of generations after the intervention, and was proportional to connectivity rate. For example, 60 generations after the intervention the fitness benefit in population 1 (warmest population) was 57-times higher at 33% connectivity rate than at 1% connectivity rate (Supplementary Fig. 11).

#### Environmental tolerance

Environmental tolerance was measured by the standard deviation of the fitness function. Genetic adaptation was slower at higher levels of environmental tolerance (Supplementary Fig. 8) because selection pressure was mitigated by the wider envelope of environmental conditions tolerated by individuals. Notwithstanding, all three intervention scenarios had smaller, but longer lasting fitness effects at higher levels of environmental tolerance (Supplementary Fig. 13). For example, on average across all scenarios, the immediate fitness effect of an intervention was 3-times higher when the sd of the fitness function was 0.05 (low environmental tolerance) than when it was 0.3 (high environmental tolerance). At high environmental tolerance (sd = 0.3) the amplitude of the fitness ripple faded to below 0.01 after approximly 15 and 17 generations later compared to low environmental tolerance (sd = 0.05), in scenarios 1 and 2, respectively. The fitness effect of scenario 3 lasted across the entire simulation of 150 generations (only 60 generations shown on Supplementary Fig. 13.) Importantly, the fitness effect was increasingly positive even in suboptimal ENSO-like phases as environmental tolerance increased. For example, the fitness effect of intervention scenario 3 on population 4 was positive in 54% of the first 60 generations after intervention when environmental tolerance was 0.05 (i.e. 2.5% of the maximum theoretical phenotypic trait value); while it was positive in 100% of generations when environmental tolerance was 0.3 (i.e. 15% of the maximum theoretical phenotypic trait value; Supplementary Fig. 13).

**Mutation rate and the size of the receiving population** had little influence on the outcomes of the three intervention scenarios relative to the other factors tested (Fig. 2). However, two non-biological parameters were highly influential: **the ratio of inoculum size to population size** (0.06 ± 0.01), and **the timing of interventions relative to ENSO-like oscillations** (0.05 ± 0.005; Fig. 10). Higher inoculation ratios resulted in larger and longer lasting fitness effects. An increase in inoculation ratio from 10%to 80% increased fitness effects by 0.12 ± 0.02 units in the first generation after intervention across all scenarios. The amplitude of the fitness ripple faded to below 0.01 units at about 8 and 30 generations later with an 80% inoculation rate compared to a 10% rate in intervention scenarios 1 and 2, respectively. The fitness effect of scenario 3 lasted across the entire simulation of 150 generations, with increasing amplitude of oscillations (60 generations shown in Supplementary Fig. 14).

ENSO-like oscillations were incorporated into our simulations to account for the temporal variability in climate (Fig. 3A). The timing of the intervention relative to these environmental cycles proved critical for the short and long-term effects of interventions on the mean fitness of the populations. In scenarios 1 and 2 that used only existing genetic diversity for inoculation, the fitness effect of interventions was highly transitional, lasting typically less than ten generations post intervention, with a rapidly decreasing amplitude (i.e. ‘fitness ripple’; Fig. 3B; Supplementary Figs. 9-14). Due to the oscillation of the environmental optimum, the first generations following the intervention experienced either an increase or a decrease in fitness compared to no intervention (Fig. 3B). The combination of the rapid decay of the fitness ripple and the associated negative or positive fitness effects demonstrated that the timing of the intervention relative to the ENSO-like cycles critically determines whether the net fitness effect of the intervention is negative or positive over the timeframes tested (Fig. 3C). For example, to increase fitness effects, the optimal intervention time was an ENSO-neutral generation followed by an ENSO-positive period (ENSO-like phase 2 on Fig. 3). Under all three scenarios, interventions at this ENSO-like phase resulted in a positive net fitness effect in the target population in the first 20 generations after warming (Fig. 3C). Additionally, the cumulative fitness benefit of interventions at this ENSO-like phase were 5.6- and 3.4-times higher for scenario 1 and 2, respectively, than when interventions occurred in an ENSO-positive phase that was followed by an ENSO-negative period (ENSO-like phase 4 on Fig. 3; Supplementary Fig. 12). Importantly, the net effect of intervention scenario 3 (the introduction of novel genetic variation) was negative between 20 to 50 generations post-intervention even at the optimal timing; and at all other intervention times it had a net negative effect for at least 45 generations (Fig. 3C), with the exception when diversity was low (He < 0.2; Supplementary Fig. 10), environmental tolerance was high (sd of fitness function > 0.1; Supplementary Fig. 11) or the trait was encoded by over 200 genes (Supplementary Fig. 12).

##### Box 1. Case study: Implications for the feasibility of genetic interventions on coral reefs

Tropical coral reefs are among the most severely impacted ecosystems by climate change due to the sensitivity of scleractinian corals to increases in ocean temperatures ^17^. The accelerating rate of reef decline over recent decades has prompted reef managers worldwide to increasingly consider novel reef restoration initiatives, including those methods incorporating the assisted evolution of the host coral and/or their associated microbial symbionts ^2,18^. Understanding the factors that influence adaptation in the wild are critically important to the assessment of the feasibility and the risks associated with applying human interventions. The evolutionary models presented here provide a baseline against which we can compare empirical data on relevant population genomic parameters of various organisms, including connectivity and genetic diversity ^19,20^ and ecological risks ^3,21,22^. We predict that genetic diversity is one of the key factors defining adaptive rates and potentials in wild populations and will influence host-directed intervention outcomes when different scenarios are considered (scenarios 1-3 outlined in main text). Based on available estimates of genetic diversity, host adaptive capacity may be high ^23^ but a formal assessment is needed to quantify adaptive potentials. Using a Web of Science search (heterozygosity AND coral, expected heterozygosity AND coral, genetic diversity AND coral (1991-2020)), we surveyed the literature encompassing 1,150 research articles that included 51 coral species (potentially including cryptic species) from 17 coral genera across six oceanic regions. We then derived genetic diversity estimates from the alloenzyme, microsatellite and Single Nucleotide Polymorphisms (SNP) sequencing data presented in these papers and averaged by coral family and reef region. We found global estimates of genetic diversity in corals varied extensively by species, genus, and region (inset Figure). Caribbean acroporids exhibited particularly low genetic diversity, perhaps from the steep population declines that they experienced in the last few decades, although promisingly, some exceptions exist (e.g. Curacao and Guadeloupe). *Galaxea* (n=20) recorded the highest overall diversity (He= 0.85 ± 0.01 SE) and *Siderastrea* (n=15) the lowest (0.14 ± 0.01). Well-studied families like Acroporidae (n=345) exhibited intermediate diversity estimates (0.475 ± 0.01). Of the 46 reef systems for which heterozygosity of corals was calculated, the Great Barrier Reef (GBR, n=167) ranked 35^th^ (0.35 ± 0.02) in diversity, with reefs in Cuba, Venezuela, and Singapore having the highest per-reef diversity estimates (0.81-0.79 ± 0.1-0.001), and reefs in Florida, Brazil, and the Dominican Republic having the lowest (0.17-0.18 ± 0.03-0.08). Overall, global estimates of genetic diversity were higher compared to the averaged modelled values of the receiving populations (~0.4 vs ~0.18), which may result in discrepancies between modelled and empirical results.

Genetic diversity appears to be highly variable in nature, suggesting that the three intervention scenarios modelled here will have varying levels of impact on reefs globally. For example, coral populations that are relatively depleted of genetic diversity may respond quickly and positively to the influx of new genetic diversity (e.g., *Acropora cervicornis* in Florida (n=11, 0.01 ± 0.001)) while those that still have high genetic variation (i.e., He > 0.4, e.g., some Acroporidae, *Galaxea, Platygyra, Pocillopora*), may respond slower, or with little effect.

Empirical results indicate that interventions like the selective breeding of corals sourced from across the GBR significantly increase novel genetic diversity in genomic regions important for the acquisition of heat tolerance ^24^. Although encouraging, due to the costs and risks associated with human interventions, it is important that future models are based on empirical biological and environmental data explicit for specific populations targeted for intervention. This requires that further experimental studies, like those currently underway ^20,24–27^ continue to fill in the knowledge gaps on the genomic underpinnings of traits related to climate change adaptation in corals.

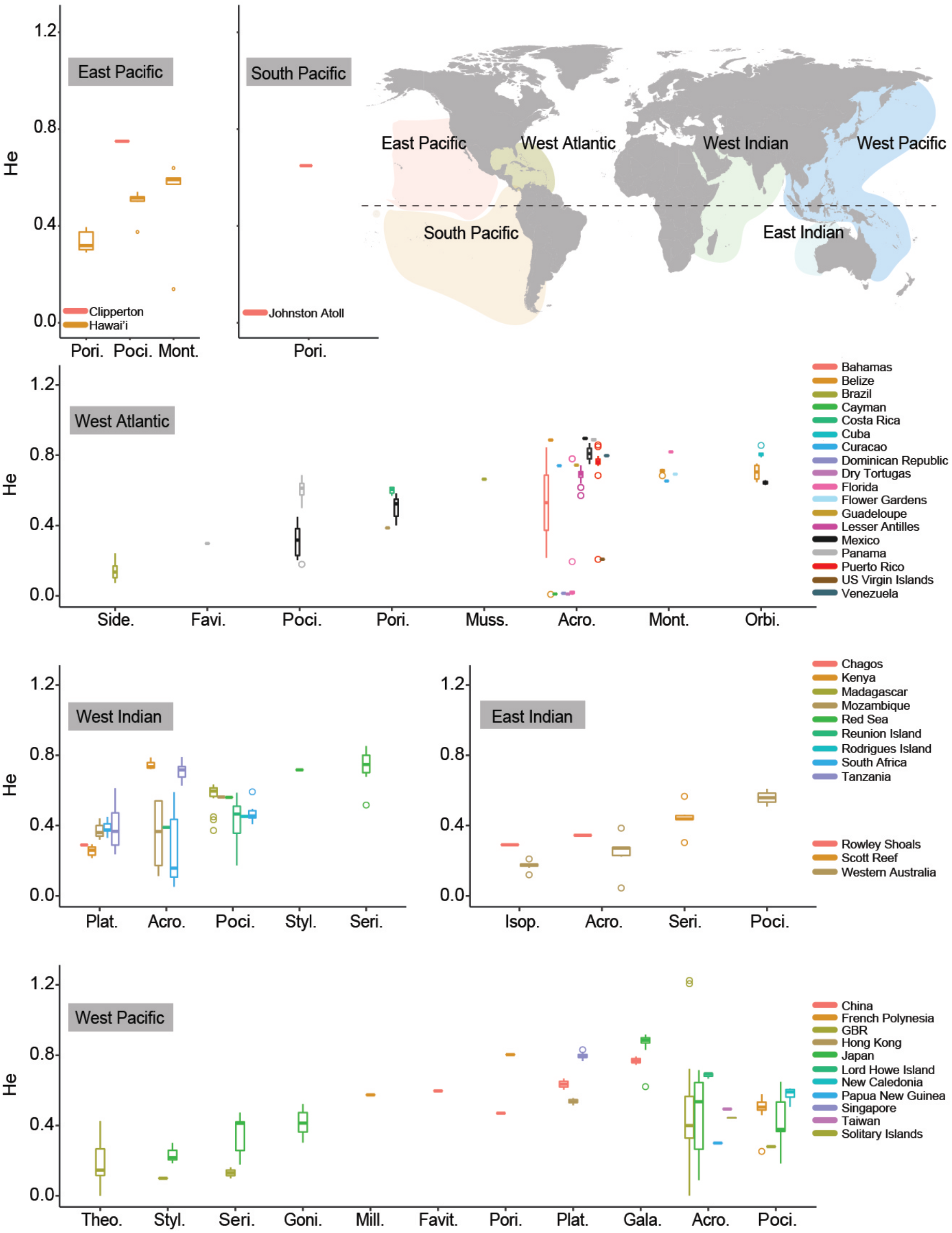

## Discussion

We demonstrate that the outcome of the modelled genetic interventions on the fitness trajectories of wild target populations is tightly linked to their natural adaptive capacity. Of the parameters measured, genetic diversity, mutation effect size, the number of QTLs, connectivity and environmental tolerance were highly influential in shaping adaptive capacities, whereas mutation rates and population size were less influential. These parameters also critically determine whether an intervention will have a fitness effect, how many generations that effect is expected to last, and whether the effect is predominantly positive or negative.

### Implications for interventions

Our models underscore that genetic diversity is a key factor in shaping the rate and scope of adaptation in populations, and quantify the range of genetic diversity values that are important for intervention feasibility in wild receiving populations. Overall, genetic diversity strongly influenced the outcome of these three modelled interventions. All modelled intervention scenarios influenced population fitness, but their effect varied by the type of intervention and by the environment of the receiving population. For example, when genetic diversity was low (He = 0 to 0.06), interventions that used only standing genetic variation (scenarios 1 and 2) had little fitness effect whilst the addition of novel diversity (scenario 3) had a strong and immediate positive fitness effect. Under higher levels of diversity (>0.2) in the receiving population, the relative influence of introducing novel diversity (scenario 3) on fitness was less. Amongst the three interventions tested, scenario 3 had the strongest impact on fitness across all diversity parameters, which was greatest and most immediate when applied to low diversity populations. In summary, novel beneficial diversity can have a large effect on evolutionary trajectories of genetically eroded populations. This is consistent with the theory and practice of genetic rescue ^28^, and highlights that genetic intervention in a population suffering from genetic erosion will likely respond differently compared to a population with higher genetic diversity. This result underscores the importance of developing an evidence-based management strategy (sensu ^9^) in which one of the first steps is to understand baseline values across key ecological and evolutionary parameters for populations and species that may be targeted for interventions.

Of the three types of interventions modelled here, the most effective are those that introduce novel genetic variation into target populations. These can either be in the form of genetic variants present in one population but not others (private alleles) potentially by reproductively crossing gene pools that are otherwise separated via physical, temporal, or reproductive barriers. Among the modelled scenarios, only the addition of novel genetic variation increased the overall adaptive potential of a population (i.e., a change beyond their original phenotypic range) and had lasting effects beyond the short term across the entire simulation of 150 generations (Supplementary Figs. 10-12). When persistent genetic changes to populations are desired, for example if warming trajectories do not appear to stop, the addition of novel genetic variation to a population would provide the greatest improvement, although it is important to note that this intervention did result in significant negative fitness consequences for ~ 40 generations under these model conditions (Fig. 3C). Alternatively, interventions that harness standing genetic variation bring about a shorter-term effect of a few generations, which may be ideal under specific management scenarios where interventions are designed to be effective over the short-term (i.e. stop-gap) between climate action and a decrease in atmospheric warming. This highlights the importance of assessing both the long-term and short-term consequences of interventions on ecosystems for both positive and negative outcomes. In summary, the variability in fitness outcomes depending on time-frames measured underscores the importance of defining the success of interventions for different stakeholder groups, in which managers may prefer to see a positive increase in fitness over shorter timescales (5 years), whereas ecologists are more concerned with positive fitness effects over longer timescales (>150 years) and highlights the need for modelling across a spectrum of temporal scales.

The timing and spatial positioning of interventions is also critical to shaping intervention outcomes. For example, shuffling the standing genetic diversity will have the strongest effect on the fitness of populations when: 1) populations are maladapted, 2) when the pace of adaptation lags behind the rate of environmental change, or 3) when inoculation happens during unfavourable phases of the ENSO-like oscillation (Supplementary Figs. 10 and 11). Interventions that do not add novel genetic variation to the metapopulation (scenarios 1 and 2) have shorter fitness effects, typically less than ten generations, and these incorporate both fitness benefits and deficits depending on the phase of a temporally variable environment (‘fitness ripples’; Fig. 3; Supplementary Figs. 10 - 14). The timing of interventions in relation to environmental cycles (e.g. ENSO) is therefore critically important in achieving an overall fitness benefit over the lifespan of the fitness ripples (Fig. 3). For example, increasing the mean temperature optimum of a population in an unusually warm year increases the fitness, but the same intervention in a colder than average year will have negative fitness effects temporarily (Fig. 3). In practical terms, interventions should be implemented in phase 2 of the ENSO cycle (Fig. 3A), from a neutral to warm period, to result in positive fitness benefit in both the short and longer term. Ultimately, it is important to also consider the ecological consequences of both the positive and negative fitness effects caused in the various phases of the environmental oscillation.

Our modelled interventions focussed on shifting the mean of the fitness curve (Fig. 5) and quantified how the breadth of phenotypic variation influences organisms’ capacity to adapt to changing environmental conditions. The extent of phenotypic variation has long been implicated as a driving force in adaptation ^29^ and there is evidence that particular forms of assisted evolution approaches on organism like dinoflagellates, insects, and corals can influence phenotypic variation by increasing the width of the fitness function instead of the mean (Fig. 5) ^24,30,31^. These approaches are promising because a wider environmental envelope buffers against environmental change and oscillation; but they also carry potential long-term risks because environmental tolerance may also slow down adaptation by decreasing the selective pressure ^32^. In our models, for example, the receiving populations with broader environmental tolerance showed slower rates of adaptation. Theory suggests that changes in allele frequencies, and therefore adaptation rates, can either be hindered or facilitated by the extent of a populations’ environmental tolerance. For example, adaptation is generally slower if environmental tolerance is wide, because the overall selection pressure decreases as it is exerted over a wider phenotypic range. Simultaneously, adaption rates may increase under wide environmental tolerance because it allows populations to move into and persist in new environments, thereby promoting selection. This example highlights the interplay between eco-evolutionary factors (reviewed in ^33^) and underscores the importance of developing eco-evolutionary models that are able to quantitatively assess complex interactions and be tailored to species and intervention strategy to predict the impacts of either taking (or not taking) specific management actions.

Overall, our models provide guidelines for future work supporting strategic management decisions and highlight that interventions should be tailored to the level of standing genetic diversity (albeit in conjunction with other relevant factors). The threshold value of He < 0.2 calculated from our models indicates a level of genetic diversity above which intervention is not warranted or may likely have little benefit. Using this threshold value in conjunction with results of our meta-analysis on genetic diversity in coral populations (see Box 1) can help guide managers through assessing the risks, costs and benefits of conservation actions (Anthony et al. 2020). It is also important to note that the connection between genetic diversity and population health is complex, where “low” values may not be indicative of degraded systems (i.e. Lewontin’s Paradox). Even ecosystems generally characterized as near pristine (e.g. the GBR) averaged genetic diversity values of He = 0.35. The value of He < 0.2 is merely a threshold of greatest intervention impact, but whether ecosystems should be allowed to potentially degrade or shift to that degraded state is left to be determined through the incorporation of other decision-science metrics. We acknowledge that other considerations exist, for example, the loss of specific ecosystem services or the loss in socio-cultural values, which could make the threshold for when to intervene increase to greater He values.

Finally, the levels of environmental tolerance and environmental heterogeneity, both present and projected for the future, should also be considered in specific management decisions. Our results suggest that the greatest positive impact is expected from interventions that introduce novel genetic diversity (e.g. scenario 3), and the timing related to environmental oscillations is critical. For example, if the inoculum consists of individuals that have been bred for increased heat tolerance only (i.e. directional selection, where the mean of the fitness curve is shifted in one direction; ^9^, the best fitness outcome is achieved by an intervention timed at the end of a cold ENSO-like phase heading into a warmer one. To ameloriate the effect of a fitness deficit during cold ENSO cycles, inoculum can be bred for increased environmental tolerance (i.e, the widening of the fitness function). Encouraginly in species like corals, some first generation interpopulational hybrids from specific reef by reef reproductive crosses did not suffer decreased fitness (as defined by both survival and growth) when outplanted to the field in a cold year ^34^.

All models balance generality, precision, and realism ^35^ and the models presented here are by design general, applicable to a diversity of life-histories and are not tailored to any specific organism (e.g. R-vs. K-strategists, mobile or sessile, etc.). We chose to explore the importance of population genomic parameters for genetic intervention practices using Wright-Fisher models because they are relatively simple and general, have an extensive history, and are relatable to previous studies (e.g. ^19,20^). The goal of this paper was to demonstrate and clarify the principles of selection on monogenic and polygenic traits in a variety of intervention scenarios through the iteration of population genomic parameters suspected as being important for genetic interventions, but yet untested in models or in practice. We confirm that genetic diversity and the number of loci determining a trait have the greatest effect on the rate and scale of adaptation; and perhaps more importantly, we quantify the parameter space in which these variables become important for maximizing intervention effectiveness. However, we recognize that for specific management decisions, even more targeted models are needed. It is increasingly evident, that most traits have high levels of underlying genetic variation in nature, and that the response to selection is often constrained by both genetic diversity and by interactions amongst genes as well as amongst traits ^7,36^. Therefore, a detailed understanding of the genetic architecture of fitness related traits, including epistasis and linkage maps; as well as the trade-offs among traits is critical for improved evolutionary modelling efforts in the future. Additionally, and especially important on coral reefs, the contribution of microbial partners to the phenotypes of their hosts is increasingly recognised ^37^, and the interplay between the evolution of the host organism and its microbial associates cannot be neglected in coral adaptation models ^23,25,38,39^.

Finally, the ecological characteristics of species (e.g. life histories, intra- and interspecific interactions) profoundly influence adaptive processes via eco-evolutionary feedback loops ^10^. Threshold values for key ecological and evolutionary parameters are sorely needed to guide the development of a portfolio of active interventions on ecosystems, like corals reefs, that are under extreme pressure from anthropogenic stressors. A genomic modelling framework, like the one provided here, supports the risk/benefit assessments and decisions to be undertaken, including whether deployment is needed, and if it is, where, when, and what to deploy. It is important to note that the predictive power of these models should be further strengthened by parameterizing with spatially explicit, taxon-specific empirical data, fine-scale empirical and modelled predictions on the rate of environmental change (e.g. for coral reefs see ^40^), and simulating more realistic non-Wright-Fischer populations. In summary, this paper provides key threshold values, and a generalized framework of how genomic models can guide decision support around active interventions and underscores the importance of the rapid development of genomic and ecological modelling approaches in target species for conservation and restoration.

## Methods

### SLiM modelling

To illustrate the principles of genetic adaptation and the effect of genetic interventions for monogenic vs. polygenic traits in isolated populations vs. in metapopulations, we simulated the evolution of Wright-Fischer (WF) populations using SLiM 3.3 ^8^. WF population models are useful to demonstrate the principles of evolutionary processes due to their relative simplicity and have been used widely in previous modelling efforts to understand the evolutionary trajectories of coral populations (e.g. ^19,20^). Starting from simple, single-gene, single-population models, working towards more complexity, we demonstrate the fate of introduced alleles and genotypes, and their impact on the fitness trajectories of the recipient populations or metapopulations. The models are built in a hierarchically complex manner, i.e. each model builds on the machinery of the previous one, for easier cross comparison. The simpler models simulate textbook evolutionary scenarios and can be found in Supplementary material. Here we only present the most complex metapopulation model on which we explored the parameter space for a number of key population genomic attributes to assess their importance for model outcomes.

#### The metapopulation and its environment

Five populations were arranged in a stepping stone metapopulation along an environmental (e.g. temperature) gradient, with population 1 having the highest phenotypic optimum trait value (1.1), and population 5 the lowest (0.9; Fig. 4). These values can be easily rescaled, see below. The environmental gradient can be thought of as a temperature gradient along latitudes, and the phenotypic trait under selection as temperature tolerance, therefore, population 1 is warm-adapted, and population 5 is cold adapted. The environment was kept stable for 100 generations (burn-in), after which an El Niño Southern Oscillation (ENSO) – like cycle with an amplitude of 0.1 phenotypic trait value units, and a wavelength of five generations was faded in over 100 generations. Note that in WF models generation time is not specified, therefore the ENSO-like oscillation cannot be considered a 5-year cycle, it is merely a proxy for non-linear environmental variability. A warming of 0.01 phenotypic trait value per generation started at generation 200 and lasted 150 generations, to simulate global warming associated with anthropogenic climate change.

**Figure 4.**
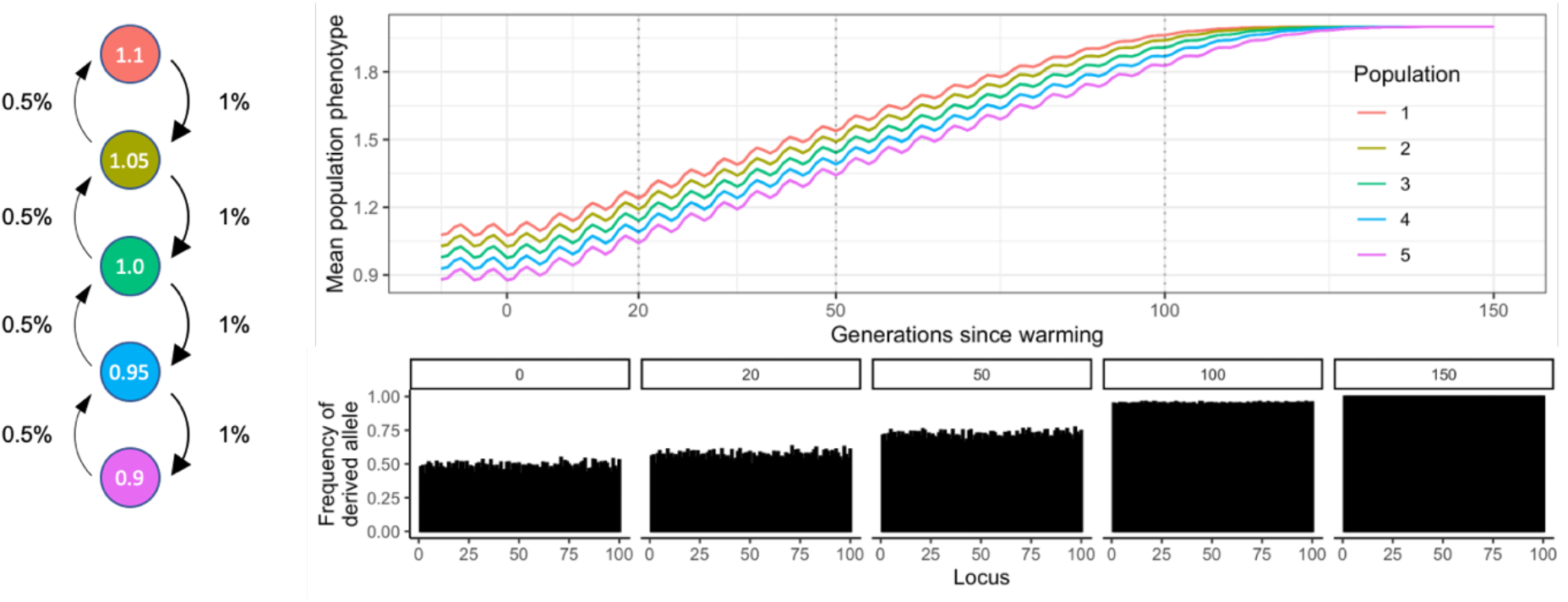
Diagram of metapopulation structure including migratation rates, environmental conditions and allele frequencies at sampling timepoints of the SLiM model discussed in this study.

Our simulations employed a mechanistic approach of additive genetic variance to estimating quantitative trait loci (QTL)-based phenotypes and fitness. Phenotypes of individuals were calculated as the sum of the effect size of all derived alleles present in an individual at each QTL (Fig. 5). The maximum phenotypic trait value was therefore maximized when an individual was homozygous for all derived alleles with a positive effect, and had no derived alleles with a negative effect. For simplicity, in this study mutation effects were kept uniformly at +0.01 except when explicitly testing for the influence of mutation effect sizes on adaptive processes. Therefore, an individual homozygous at 100 QTLs had a phenotypic trait value of 2.0 (100 QTLs * 0.01 mutation effect * 2 chromosomes).

**Figure 5.**
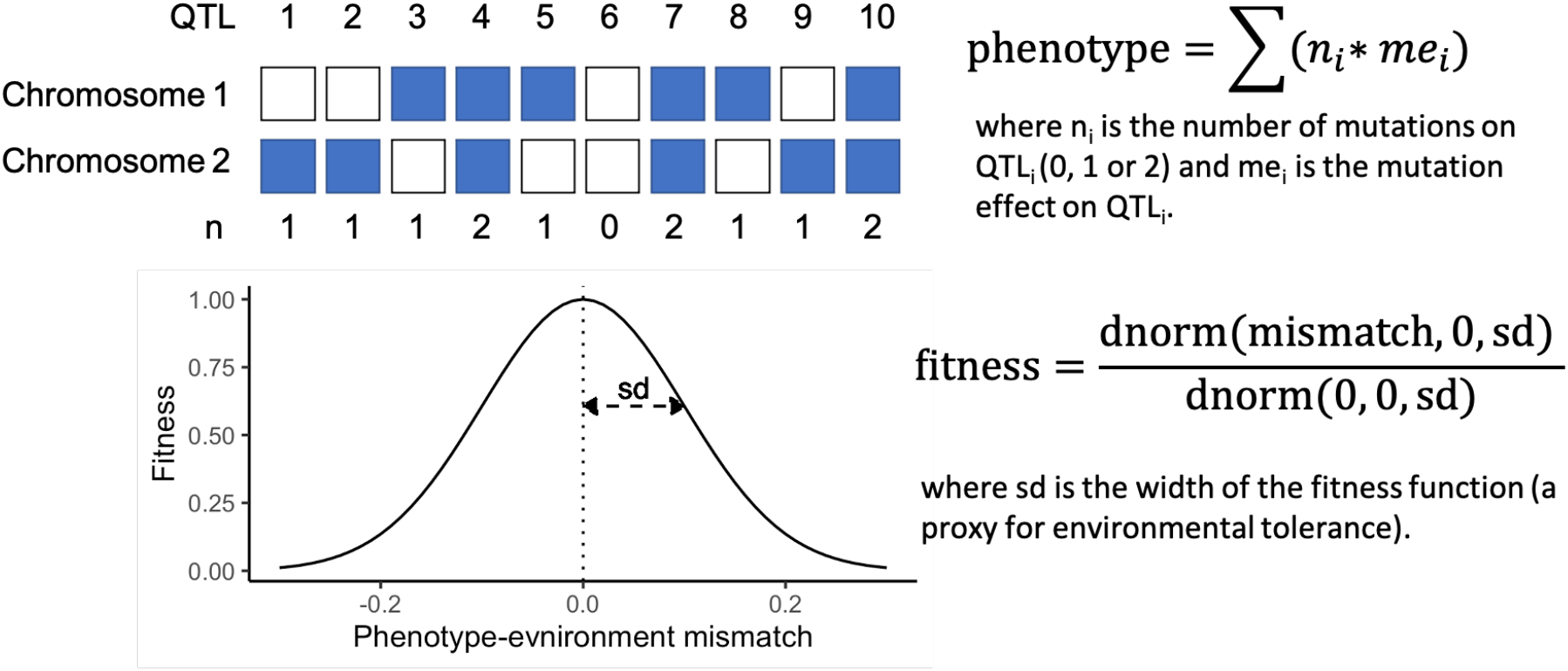
A mechanistic approach to QTL-based phenotype and fitness calculations, with a schematic for 10 QTLs.

Individual fitness was calculated from the mismatch between each individual’s phenotype and its environment, following a Gaussian density function with a mean of 0 and standard deviation (sd) of 0.1 (except when testing for the effect of environmental tolerance on adaptive processes; Fig. 5). The width of the fitness curve, defined by the sd of the Gaussian density function, was used as a proxy for environmental tolerance. Note that phenotypic trait values, environmental optima, rate of warming, the amplitude of ENSO-like cycles, mutation effect and the width of the fitness function are meaningful only relative to each other. The values chosen for this study can be easily rescaled. For example, a model with ten times higher values for these six parameters (i.e, warming = 0.1°C per generation, environmental optima = 9 – 11°C, amplitude of ENSO-like cycles = 1°C, mutation effect = 0.1, and sd of the Gaussian fitness function = 1.0) will yield the same results as the values described above. Furthermore, the values of environmental optima can be re-centred freely, therefore the above model describes the behaviour of a metapopulation that populates any temperature gradient with 2 degrees difference between the warm and cold extremes, which is in the range for many natural systems, e.g. corals on the Great Barrier Reef (Quigley et al 2020).

It follows from this mechanistic approach to estimating phenotype and fitness that the adaptive range of a population is defined by the number of QTLs and their respective mutation effects, because these constrain the range of theoretically possible phenotypic trait values (maximum – minimum phenotypic trait value). If the environmental optimum increases over time (e.g. simulated global warming), the frequencies of alleles with a positive fitness effect shift towards fixation, while alleles with a negative fitness effect are gradually lost. As a consequence, genetic diversity (heterozygosity) decreases, and the metapopulation starts saturating its adaptive range. When all QTLs fix or are lost, populations can no longer evolve without the addition of novel genetic diversity via gene flow, mutation, or active human intervention (Fig. 4).

#### Model parameters and their influence on adaptive processes

The parameter space and relative importance of each parameter of the model was explored by varying one parameter at a time while keeping the others constant. The relative importance of each model parameter was expressed as the standard deviation of the model output (e.g. mean population fitness at a certain generation) under the group of parameter values tested. We explored the importance of the following model parameters for the outcome of simulations:

##### Connectivity

Connectivity is expressed as the proportion of offspring in a population produced by parents of another population. Our metapopulation model used a stepping-stone structure, where each population was only connected to an upstream and a downstream population directly. Connectivity was twice as high downstream than upstream, and population 1 was the most upstream population, while population 5 the most downstream (Fig. 4). The range of values explored to test the influence of connectivity on adaptive processes were 33% to 0.001% migration downstream; and the value at which other parameters were tested was 1% downstream connectivity.

##### Genetic diversity

Genetic diversity is a key metric in influencing adaptation rates. It is also critically important in conservation given that genetic erosion can lead to inbreeding depression and an increase in extinction risk through the erosion of evolutionary potential. Genetic diversity is measured in a variety of ways, including allelic richness (number of alleles per locus), observed and expected heterozygosity (e.g. Nei’s unbiased measure, He ^41^), haplotype diversity (h), nucleotide diversity (π ^42^), and gene diversity (H ^43^). Effective population size (*Ne*) has also been used as a proxy for genetic diversity and direct measure of adaptive potential ^19,44,45^. Expected heterozygosity is a widely cited metric of diversity which describes the number of heterozygous loci in populations, and is therefore of critical importance in quantifying the capacity for populations to cope with environmental change (see above; Fig. 4) and can be manipulated directly in the SLiM models. In our models, genetic diversity was measured as heterozygosity (He) across all loci. To achieve various levels of He, we directly manipulated the number of QTLs while keeping mutation effect size constant. When mutation effect is kept constant, the number of QTLs in the model defines the maximum phenotypic trait value (MPTV) achievable in the population. As the environmental optimum gets closer to the MPTV, QTLs fix for the derived allele and genetic diversity (heterozygosity) drops (Fig. 4). Therefore, the number of QTLs in our models are directly proportional to heterozygosity. The range explored was 60 to 120 QTLs coding the trait with a uniform mutation effect of +0.01. This range translates to 1.2 and 4.0 MPTV, respectively, in an environment where the phenotypic optima of populations ranged from 0.9 to 1.1 before warming, and 2.4 to 2.6 by the end of a 150-generation warming period; which in turn translates to 0-0.48 values of heterozygosity by the time of intervention at 50 generations post-warming. The value at which other parameters were tested was He = 0.4 (MPTV = 2.0, 100 loci).

##### Number of genes

To explore the influence of the number of genes under selection on adaptive processes, we directly and simultaneously manipulated mutation effect size and the number of QTLs, in an inversely proportional manner. This way the maximum phenotypic trait value remained constant and hence heterozygosity was not affected. For example, 10 QTLs with a mutation effect of 0.1 have the same MPTV as 100 QTLs with a mutation effect of 0.01 (i.e., 2.0). Here we explored the range 50 - 1,000 loci while adjusting mutation effect using the formula *mutation effect = 1/nQTL*. Other parameters were tested at a value of 100 loci (0.01 mutation effect).

##### Mutation rate

The rate at which new derived alleles are formed on loci. The range explored was 0 (no mutation, all diversity pre-defined) to 10e-5. The value at which other parameters were tested was 0 (no mutation).

##### Environmental tolerance

The proxy used for environmental tolerance was the width of the fitness curve, i.e., the standard deviation of the Gaussian density function, as it defines the fitness drop when individuals’ phenotypes deviate from the optimum phenotype dictated by the environment (Fig. 5). The wider the fitness curve, the more tolerant individuals are to mismatching their environment. The range explored was 0.01 to 0.3, and the default value at which other parameters were tested was 0.1

##### Population size

the number of individuals in each population of the metapopulation. This value was identical in all five populations for simplicity, and kept constant over time. The range explored was from 1,000 to 20,000 individuals per population and the default value at which other parameters were tested was 10,000.

To quantify the importance of these key population genomic parameters for determining adaptive capacities of populations, we ran the simulations 10 times for each permutation of model parameters while holding the other parameters constant. For each simulation and parameter combinations, we then calculated the standard deviation of the (i) fitness change over 20 generations of warming as a proxy for the rate of adaptation; and ii) of the mean population fitness achieved by 20, 50 and 100 generations of warming as a proxy for the scope of adaptation. The higher these metrics (the standard deviation of possible model outcomes), the more influential the parameter was considered for adaptive rate and realised adaptation.

#### Intervention parameters and their influence on the outcome of interventions

Given the parameters described above, three scenarios where simulated to model the effects of active genetic interventions on the fitness trajectory of wild populations. We added individuals to the metapopulation that were created by i) crossing individuals from the extreme populations (population 1 and 5; simulating assisted gene flow; scenario 1); ii) artificially reshuffling the standing genetic variation of the metapopulation, so that the inoculum had a phenotype that perfectly matched the phenotypic optimum of the recipient population (simulating selective breeding; scenario 2); or iii) adding new alleles (novel genetic variation) to the existing genetic variation, so that the inoculum had a phenotype that perfectly matched the phenotypic optimum of the recipient population (simulating assisted translocation or direct genetic modifications; scenario 3). Inoculum was added to either population 3 that lies in the middle of the environmental gradient (hence with an average environment); or to population 4, that is slightly cooler than the average.

Additional to the above population genomic parameters, we explored the importance of the following intervention-specific model parameters on the outcome of interventions:

##### Inoculation ratio

the proportion of newly introduced individuals (inoculum) to the size of the recipient population. The range explored was 1% to 80%, and the default value at which other parameters were tested was 50%.

##### Timing of intervention

to test the effect of the timing of intervention in relation to ENSO-like environmental oscillations, we simulated interventions in five consecutive generations that represent different ENSO-like phases (Fig. 3). Note that because warming was simultaneously ongoing, a slight warming signal was unavoidable in these comparisons.

##### Brood stock

the number of parent colonies used in lab-based crossing of the extreme populations. The range explored was 50 to 1,000 colonies, with 100 colonies as a default value at which other parameters were tested.

The relative importance of intervention parameters and population genomic parameters on the fitness trajectories following interventions was quantified by the sd of the fitness difference between control (no intervention) and treatment (one of three intervention scenarios) simulations, for each parameter while holding the other parameters constant. The larger the sd of the Δ fitness between control and treatment, the more influential the parameter was considered for the development of fitness trajectories.

## Code availability

SLiM model code, output data, and R scrips and markdowns to graph and analyse the data are available at https://github.com/dontorda/ and https://github.com/LaserKate.

## Acknowledgments

The authors would like to thank Orpheus Island Research Station staff for their support during the writing workshop, where this manuscript was developed. GT was funded by the Australian Research Council’s Discovery Early Career Researcher Award (DECRA DE200101064). KQ was funded by the Australian Institute of Marine Science.

## References

1. Scheffer, M., Carpenter, S., Foley, J. A., Folke, C. & Walker, B. Catastrophic shifts in ecosystems. Nature 413, 591–596 (2001).

2. van Oppen, M. J. H., Oliver, J. K., Putnam, H. M. & Gates, R. D. Building coral reef resilience through assisted evolution. Proc Natl Acad Sci USA 112, 2307–2313 (2015).

3. Thiele, L. P. Nature 4.0: Assisted evolution, de-extinction, and ecological restoration technologies. Global Environmental Politics 20, 9–27 (2020).

4. Fancourt, B. A. Diagnosing species decline: a contextual review of threats, causes and future directions for management and conservation of the eastern quoll. Wildl. Res. 43, 197–211 (2016).

5. Reusch, T. B. H. & Wood, T. E. Molecular ecology of global change. Molecular Ecology 16, 3973–3992 (2007).

6. Franks, S. J. & Hoffmann, A. A. Genetics of climate change adaptation. Annual Review of Genetics 46, 185–208 (2012).

7. Walsh, B. & Blows, M. W. Abundant genetic variation + strong selection = multivariate genetic constraints: a geometric view of adaptation. Annu. Rev. Ecol. Evol. Syst. 40, 41–59 (2009).

8. Haller, B. C. & Messer, P. W. SLiM 3: Forward genetic simulations beyond the Wright–Fisher model. Mol Biol Evol 36, 632–637 (2019).

9. Gaitán-Espitia, J. D. & Hobday, A. J. Evolutionary principles and genetic considerations for guiding conservation interventions under climate change. Global Change Biology 27, 475–488 (2021).

10. Hendry, A. P. Eco-evolutionary dynamics. (Princeton University Press, 2016).

11. Campbell-Staton, S. C. et al. Winter storms drive rapid phenotypic, regulatory, and genomic shifts in the green anole lizard. Science 357, 495–498 (2017).

12. Johnson, W. E. et al. Genetic restoration of the Florida Panther. Science 329, 1641–1645 (2010).

13. Hasselgren, M. et al. Genetic rescue in an inbred Arctic fox (Vulpes lagopus) population. Proceedings of the Royal Society B: Biological Sciences 285, 20172814 (2018).

14. Weeks, A. R. et al. Genetic rescue increases fitness and aids rapid recovery of an endangered marsupial population. Nat Commun 8, 1–6 (2017).

15. Hardisty, P., Roth, C., Silvey, P., Mead, D. & Anthony, K. R. N. Reef Restoration and Adaptation Program – Investment case. A report provided to the Australian Government from the Reef Restoration and Adaptation Program. 100 (2019).

16. National Academies of Sciences, Engineering, and Medicine. A research review of interventions to increase the persistence and resilience of coral reefs. (National Academies Press, 2019).

17. Hughes, T. P. et al. Spatial and temporal patterns of mass bleaching of corals in the Anthropocene. Science 359, 80–83 (2018).

18. Anthony, K. et al. New interventions are needed to save coral reefs. Nature Ecology & Evolution 1, 1420–1422 (2017).

19. Matz, M. V., Treml, E. A., Aglyamova, G. V. & Bay, L. K. Potential and limits for rapid genetic adaptation to warming in a Great Barrier Reef coral. PLOS Genetics 14, e1007220 (2018).

20. Quigley, K. M., Bay, L. K. & van Oppen, M. J. H. The active spread of adaptive variation for reef resilience. Ecology and Evolution 9, 11122–11135 (2019).

21. van Oppen, M. J. H. van et al. Shifting paradigms in restoration of the world’s coral reefs. Global Change Biology 23, 3437–3448 (2017).

22. Sandler, R. The ethics of genetic engineering and gene drives in conservation. Conservation Biology 34, 378–385 (2020).

23. van Oppen, M. J. H. & Medina, M. Coral evolutionary responses to microbial symbioses. Philosophical Transactions of the Royal Society B (2020) doi:10.1098/rstb.2019.0591.

24. Quigley, K. M., Bay, L. K. & van Oppen, M. J. H. Genome-wide SNP analysis reveals an increase in adaptive genetic variation through selective breeding of coral. Molecular Ecology 29, 2176–2188 (2020).

25. Quigley, K. M., Randall, C. J., van Oppen, M. J. H. & Bay, L. K. Assessing the role of historical temperature regime and algal symbionts on the heat tolerance of coral juveniles. Biology Open 9, bio047316 (2020).

26. Thomas, L. et al. Mechanisms of thermal tolerance in reef-building corals across a fine-grained environmental mosaic: lessons from Ofu, American Samoa. Front. Mar. Sci. 4, (2018).

27. Fuller, Z. L. et al. *Population genetics of the coral* Acropora millepora: Towards a genomic predictor of bleaching. http://biorxiv.org/lookup/doi/10.1101/867754 (2019) doi:10.1101/867754.

28. Whiteley, A. R., Fitzpatrick, S. W., Funk, W. C. & Tallmon, D. A. Genetic rescue to the rescue. Trends in Ecology & Evolution 30, 42–49 (2015).

29. Lynch, M. & Walsh, B. Genetics and analysis of quantitative traits. vol. 1 (Sinauer Sunderland, MA, 1998).

30. Quigley, K. M., Roa, C. A., Beltran, V. H., Leggat, B. & Willis, B. L. Experimental evolution of the coral algal endosymbiont, Cladocopium goreaui: lessons learnt across a decade of stress experiments to enhance coral heat tolerance. Restoration Ecology e13342 (2021) doi:https://doi.org/10.1111/rec.13342.

31. Kristensen, T. N., Loeschcke, V. & Hoffmann, A. A. Can artificially selected phenotypes influence a component of field fitness? Thermal selection and fly performance under thermal extremes. Proceedings of the Royal Society B: Biological Sciences 274, 771–778 (2007).

32. Ghalambor, C. K., McKAY, J. K., Carroll, S. P. & Reznick, D. N. Adaptive versus non-adaptive phenotypic plasticity and the potential for contemporary adaptation in new environments. Funct Ecology 21, 394–407 (2007).

33. Hendry, A. P. Key questions on the role of phenotypic plasticity in eco-evolutionary dynamics. JHERED 107, 25–41 (2016).

34. Quigley, K. et al. Variability in fitness trade-offs amongst coral juveniles with mixed genetic backgrounds held in the wild. Front. Mar. Sci. 8, (2021).

35. Levins, R. The strategy of model building in population biology. American Scientist 54, 421–431 (1966).

36. Hoffmann, A. A. & Sgrò, C. M. Climate change and evolutionary adaptation. Nature 470, 479–485 (2011).

37. Theis, K. R. et al. Getting the hologenome concept right: an eco-evolutionary framework for hosts and their microbiomes. mSystems 1, (2016).

38. Apprill, A. Marine animal microbiomes: toward understanding host–microbiome interactions in a changing ocean. Front. Mar. Sci. 4, (2017).

39. Torda, G. et al. Rapid adaptive responses to climate change in corals. Nature Clim. Change 7, 627–636 (2017).

40. Heron, S. F., Maynard, J. A., van Hooidonk, R. & Eakin, C. M. Warming trends and bleaching stress of the world’s coral reefs 1985–2012. Scientific Reports 6, 38402 (2016).

41. Nei, M. Estimation of average heterozygosity and genetic distance from a small number of individuals. Genetics 89, 583–590 (1978).

42. Huang, D., Goldberg, E. E., Chou, L. M. & Roy, K. The origin and evolution of coral species richness in a marine biodiversity hotspot. Evolution 72, 288–302 (2018).

43. Prada, C. et al. Cryptic diversity hides host and habitat specialization in a gorgonian-algal symbiosis. Molecular Ecology 23, 3330–3340 (2014).

44. Foster, N. L. et al. Connectivity of Caribbean coral populations: complementary insights from empirical and modelled gene flow. Molecular Ecology 21, 1143–1157 (2012).

45. Robitzch, V., Banguera-Hinestroza, E., Sawall, Y., Al-Sofyani, A. & Voolstra, C. R. Absence of genetic differentiation in the coral *Pocillopora verrucosa* along environmental gradients of the Saudi Arabian Red Sea. Front. Mar. Sci. 2, (2015).

