## Supplementary material for "Drivers of adaptive capacity in wild populations: implications for genetic interventions"

### From simple to complex evolutionary models of genetic interventions

To illustrate the principles of genetic interventions for monogenic vs. polygenic traits, in isolated populations vs metapopulations, we created evolutionary models of Wright-Fischer (WF) populations in the evolutionary modelling framework SLiM 3.3. Despite their disconnect from reality, WF population models are extremely useful to demonstrate the principles of evolutionary processes due to their simplicity. Importantly, WF models have been used widely in earlier modelling efforts, and we felt it was important to make our models relatable to this previous body of work. Starting from simple, working towards more complex models we demonstrate the fate of introduced alleles and haplotypes, and their impact on the fitness of the recipient populations or metapopulations. The models are built in a hierarchical manner with increasing complexity, i.e. each model builds on the machinery of the previous one, for cross comparison. The simpler models simulate textbook evolutionary scenarios. Model codes are archived on https://github.com/dontorda/

**Model 1. One gene one population**

The simplest model is a codominant single locus model of one single population. The beneficial allele 'A' has a selection coefficient of 0.1. A population of 10,000 individuals naïve to the 'A' allele receives an inoculum of 100 AA homozygous individuals. The adaptive allele spreads rapidly and fixes within 250 generations, increasing the population fitness from 1.0 to 1.1.

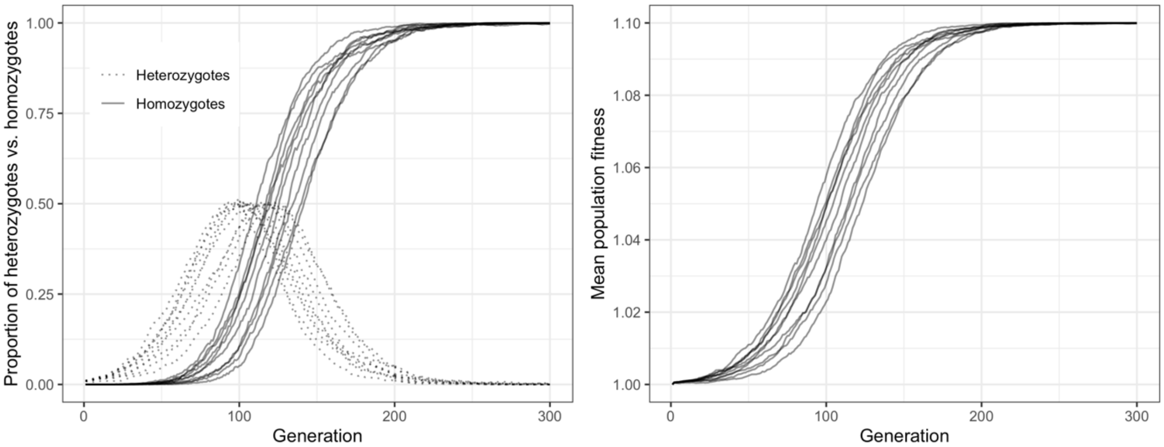


Supplementary Figure 1. Fixation of a beneficial mutation on one single locus in a population (Ten independent simulations shown).

**Model 2. One gene, metapopulation, no spatial variation in environment**

The next model is the multi-population version of Model 1. Here we have five populations with identical selection coefficient on the 'A' allele (i.e. similar environment). We assume a 1% migration rate from low to high population numbers (1 to 5), and 0.5% the opposite direction, among neighbouring populations, representing a stepping-stone metapopulation model. Each population has 10,000 individuals, homozygous for the 'a' allele, naïve for the 'A' allele. We then introduce 100 individuals that are homozygous for the beneficial allele 'A' in population 1. The allele spreads in the population and with a little lag also spreads into the other populations, increasing the mean fitness to 1.1 within 300 generations.

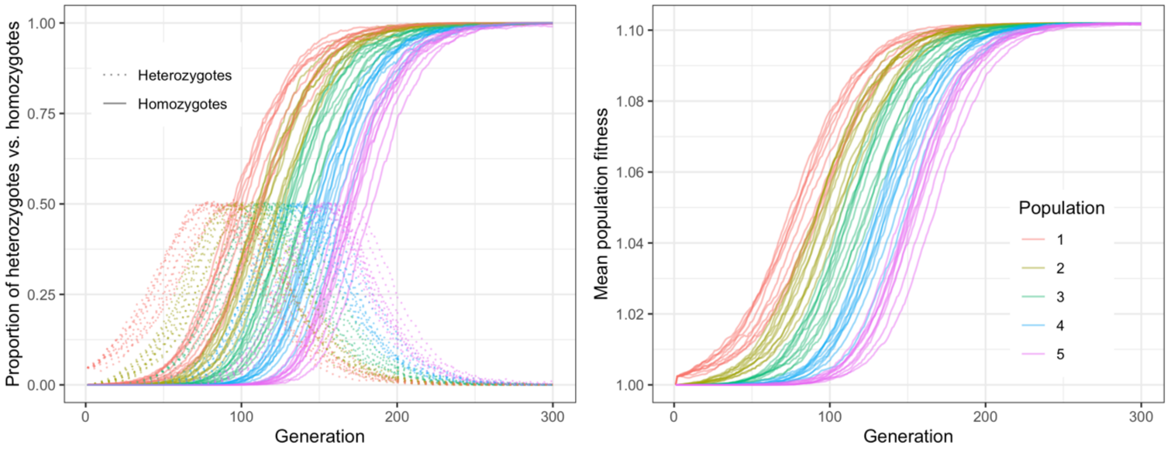


Supplementary Figure 2. Fixation of a beneficial mutation on one single locus in a metapopulation. Each population has the same selection coefficient for the new mutation. Ten independent simulations shown.

This model illustrates the basic assumption behind genetic intervention theory: the introduction of a beneficial mutation into a population of a metapopulation will create fitness benefit and spread rapidly in the system. However, it is completely unrealistic on two accounts: (i) in nature different populations have different environments, therefore the selection coefficient will vary spatially; and (ii) very few traits are coded by one single gene. Complex traits like thermal tolerance are rather coded by hundreds of loci, each with a tiny mutation effect. We will examine these factors below.

**Model 3. One gene, metapopulation, environmental variation among populations**

Building on Model 2., this time each population has a different environmental condition, representing a stepping stone model along a latitudinal gradient, for example. The mutation that was beneficial in population 1 (e.g. low latitude population), will be less beneficial as we go towards higher latitudes; in the central population (population 3) it is neutral and beyond that it is increasingly detrimental. This is a scenario that mimics the latitudinal gradient of large ecosystems, for example coral reefs, and serves as a simplified demonstration of how 'heat adapted' genotypes are excluded by environmental filtering.

Allele 'A' introduced in population 1 spreads quickly within the population, as in model 2. But contrary to model 2, the spread of the allele slows down in population 2, and even further in population 3 where the allele is neutral. In this population the allele spreads due to immigration from low latitude populations and genetic drift, but has no fitness effect. In population 4 we can observe a **strong outbreeding depression due to immigration from lower latitude populations**. Here, the 'A' allele is typically present in one copy per individual only (heterozygotes) because homozygotes are strongly maladapted and selected against. Interestingly, the magnitude of outbreeding depression is smaller in population 5 than in population 4, despite that the new allele is more detrimental there. This is because there is less intrusion of the allele via immigration from higher latitudes to this population (population 4 already acts as a filter in the stepping stone configuration).

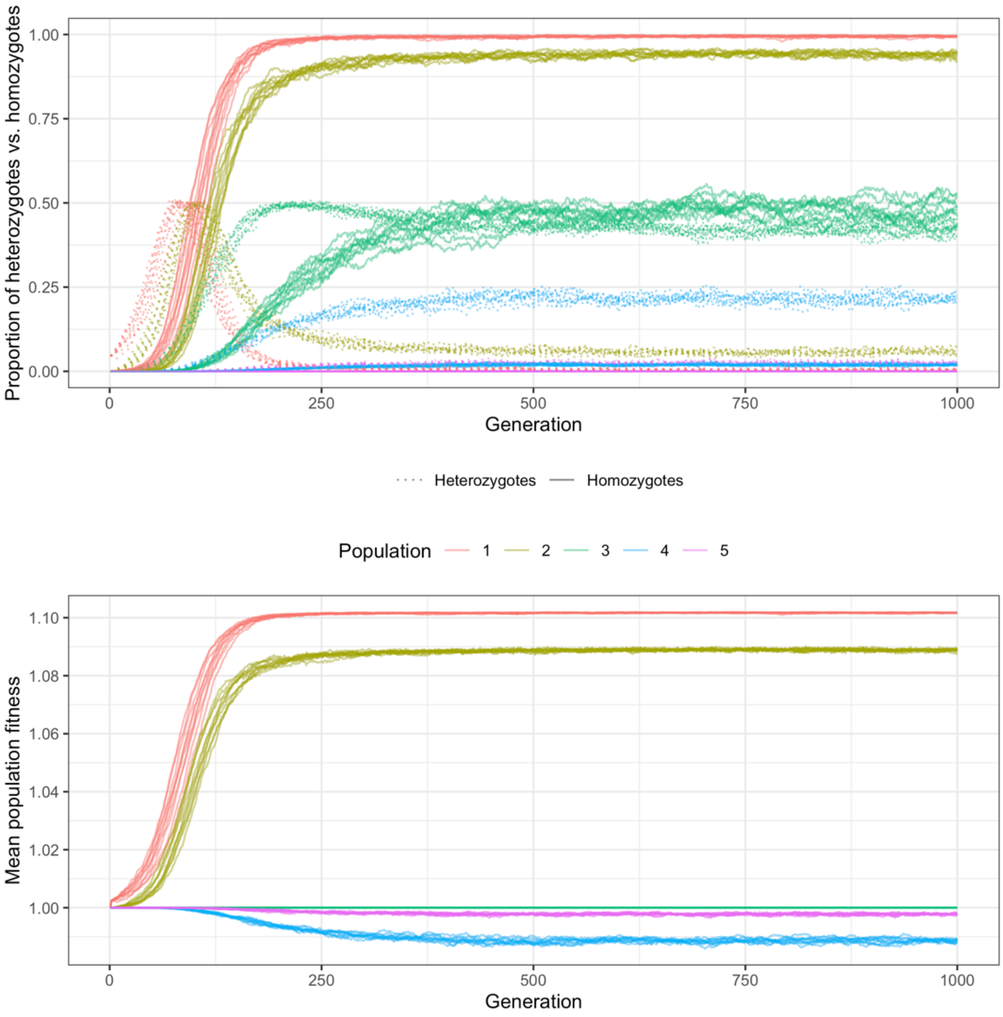


Supplementary Figure 3. Spread of a new mutation on one single locus in a metapopulation where each subpopulation has a different environment, along a cline, representing a stepping stone model along a latitudinal gradient. The mutation that was beneficial in population 1 (low latitude population), will be less beneficial as we go towards higher latitudes; in the central population (population 3) it is neutral and beyond that it is increasingly detrimental. Observe the strong fitness cost of outbreeding depression in population 4.

**Model 4. One population, polygenic adaptation**

The fourth model is once again a single population model, but this time the trait is polygenic with additive quantitative trait loci (QTLs). We explored **2 types of interventions for four completely disconnected populations** with different characteristics.

<u>The populations</u>

Each starting population was maladapted: each individual had a phenotypic trait value of 0.8 in an environment where the phenotypic optimum was 1.0. The 0.8 phenotypic value was achieved in one of two different ways:

1. In Population 1 and 2, each individual was homozygous to 40 QTLs with a mutation effect of 0.01 (40 x 2 x 0.01 = 0.8;);

2. In Population 3 and 4, each individual was heterozygous to 80 QTLS (80 x 1 x 0.01 = 0.8).

The first two populations were therefore fixed for all loci and there was no room for further evolution (one single haplotype; evolutionary dead-end). Populations 3 and 4 had a wide. adaptive range (the maximum achievable phenotypic trait value in these populations was 80 x 2 x 0.01 = 1.6) due to high genetic variation on a large number of QTLs.

<u>The interventions</u>

Populations 1 and 3 were left to evolve without intervention, whereas populations 2 and 4 were given inoculum from a lab-grown pool of perfectly adapted individuals. These optimal phenotypes were created in one of the following ways:

- Intervention 1. Individuals had 20 new QTLs that were not present in the wild population before. This is what direct genetic modification or the translocation of individuals from a completely different genetic pool would result in. These new loci, were combined with the naturally existing genetic variation to make up the inoculum that was perfectly adapted to the recipient population's environment (heterozygous to 80 natural loci and to 20 new loci = phenotypic value of 1.0);
- Intervention 2. Inoculum was created by re-shuffling only the naturally existing genetic variation on 80 loci, to create homozygotes for 20 loci and heterozygotes for 60 (hence with a phenotypic value of 1.0).

The effect of inoculum size was explored by running these interventions with 1, 10, 50 and 100% inoculum-to-recipient population ratio.

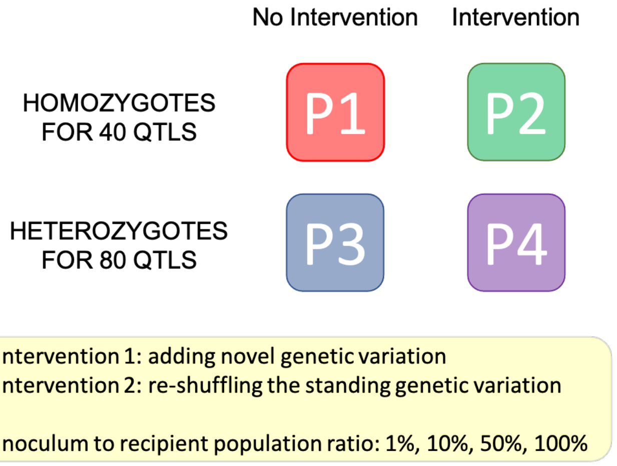


Supplementary figure 4. Schematics of Model 4.

<u>Results of Model 4</u>

Population 1 (evolutionary dead-end without intervention) remained maladapted throughout the simulation. Population 3 (good adaptive capacity, without intervention) very quickly evolved to adapt to the environmental optimum. In 10 generations the mean population phenotype climbed very close to the phenotypic optimum of 1.0. The mean fitness stagnated at around 0.9 for hundreds of generations, until the genetic variation was eroded and all individuals fixed to 50 loci and lost the rest.

Population 2 (evolutionary dead-end with intervention) benefited from the addition of new genetic variation (genetic rescue). The evolution of the mean phenotype over time was very similar between the two intervention methods (since both methods introduced new genetic variation that the population was naïve for), and was only a function of the ratio of inoculum to the recipient population. Here even a small amount of inoculum could make a big change; i.e. beneficial new variation spread quickly in the population (similar to the monogenic model, Model 1). Interestingly, mean fitness reached higher values in the first 50 generations if only small ratios of inoculum were added to the population compared to large inoculum sizes. This is because some of the introduced loci drift out immediately, and the resulting relatively lower genetic diversity creates less phenotypic variation (phenotypic variation is beneficial only in spatially or temporally heterogeneous environments). With a smaller inoculum, fewer new QTLs were added to the population, speeding up the evolutionary purging that led to achieving maximum fitness faster.

Population 4 (good adaptive capacity, with intervention) responded very similarly to the two intervention methods in terms of phenotypic change. Because it already had the adaptive capacity needed to match the phenotypic optimum (see Population 3), the addition of new alleles to the gene pool did not speed up adaptation in itself. Regardless of the actual genetic architecture of the inoculum, inoculation boosted the mean population phenotype in function of the inoculum size, and hence placed the population on an advanced trajectory towards matching the phenotypic optimum of the environment. However, the two intervention methods resulted in slightly different fitness benefits after the first 10 generations, with added genetic variation being slightly detrimental for mean population fitness for hundreds of generations compared to no-intervention or compared to reshuffling the standing genetic variation. Importantly, the fitness benefits of interventions were only significant in the first 10 generations, and only when inoculum size was large (i.e. > 50% of recipient population).

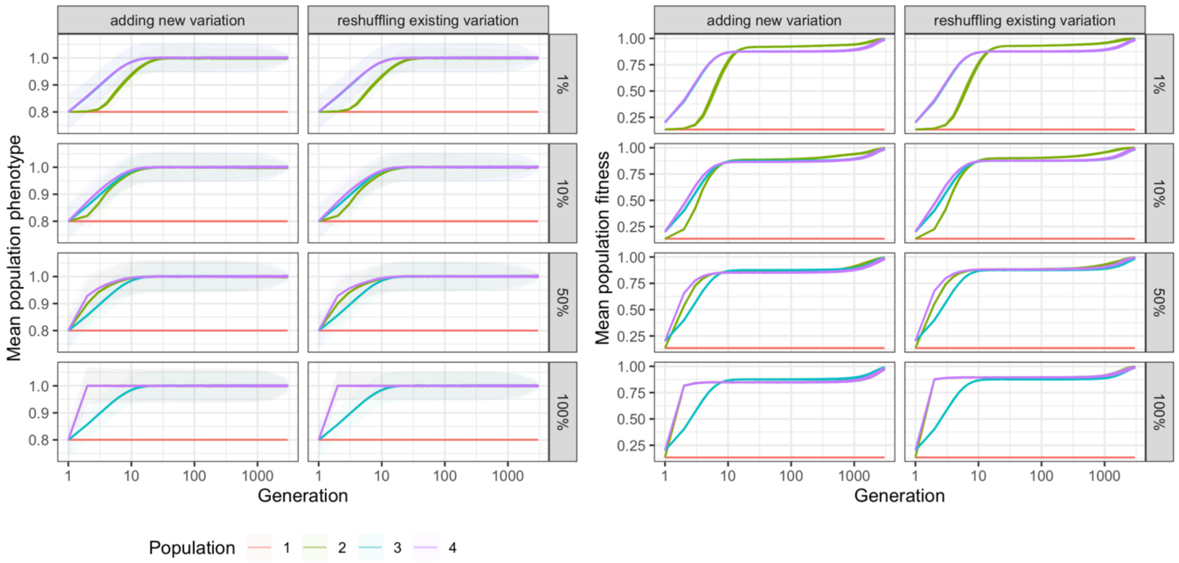


Supplementary Figure 5. Phenotype and fitness effect of two intervention types on genetically degraded (1 and 2) vs diverse (3 and 4) populations. Populations 1 and 3 were left to evolve without intervention (control), while 2 and 4 were inoculated with perfectly adapted individuals that either carried new adaptive variation, or only the locally existing genetic variants. Inoculation ratio to the recipient population is shown on facet rows.

To test the effect of genetic diversity on the efficiency of interventions, we ran Model 4 with different QTL panels, ranging from 50 to 100 loci. Because the phenotypic optimum in this model is 1.0 and the mutation effect was fixed at a uniform 0.01 value, 50 QTLs were needed for successful adaptation, while 100 QTLs provided an adaptive scope up until the 2.0 phenotypic trait value. Our simulations showed that the impact of the intervention greatly depends on the genetic diversity of the recipient population. When genetic diversity was low, even a small amount of inoculum created a large (but temporary) fitness benefit. In contrast, when genetic diversity was high, the fitness benefit was lower and faded away faster.

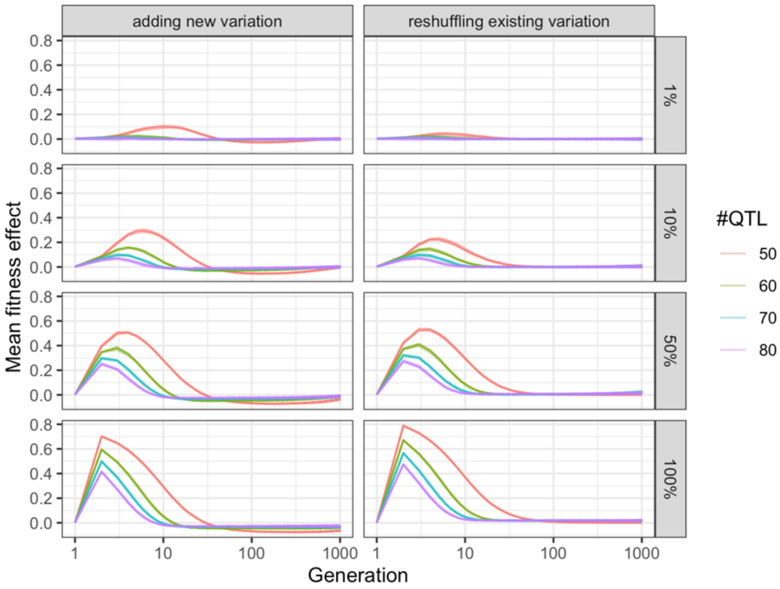


Supplementary Figure 6. Net fitness effect of two intervention types on populations with varying genetic diversity. The number of loci (#QTL, represented with colors) is directly proportional to genetic diversity since individuals strive to achieve a 1.0 phenotype with a uniform mutation effect of 0.01. When genetic diversity is low (50 QTLs, in red), even a small amount of inoculum creates a large (but temporary) fitness benefit, whereas a much larger inoculation ratio is needed to achieve the same effect in a genetically diverse population (80 QTLs, in purple). Inoculation ratio indicated in facet rows.

In summary, this model demonstrates that genetic rescue of an evolutionarily compromised, low diversity population is very different from intervening in a population with substantial genetic variation and hence adaptive capacity. A small amount of inoculum has a large effect on the evolutionary trajectory of genetically eroded populations, but large inoculum sizes are needed to reach any effect on genetically diverse populations; and the fitness benefits quickly fade away with time.

**Model 5.**

The fifth model is the final model for this manuscript, the one presented in the main body of the paper. It expands Model 4 to five populations, each with its unique environmental optimum, arranged in a stepping stone metapopulation along an environmental gradient, as in model 3. This model allows us to explore the influence of various model parameters on the rate and scope of metapopulation adaptation, and assess the fitness effect of various types of genetic interventions. For details, please refer to the Methods section of the paper. The main results are presented in the body of the manuscript, whereas here we only present some additional figures with extended captions that help understand the model results.

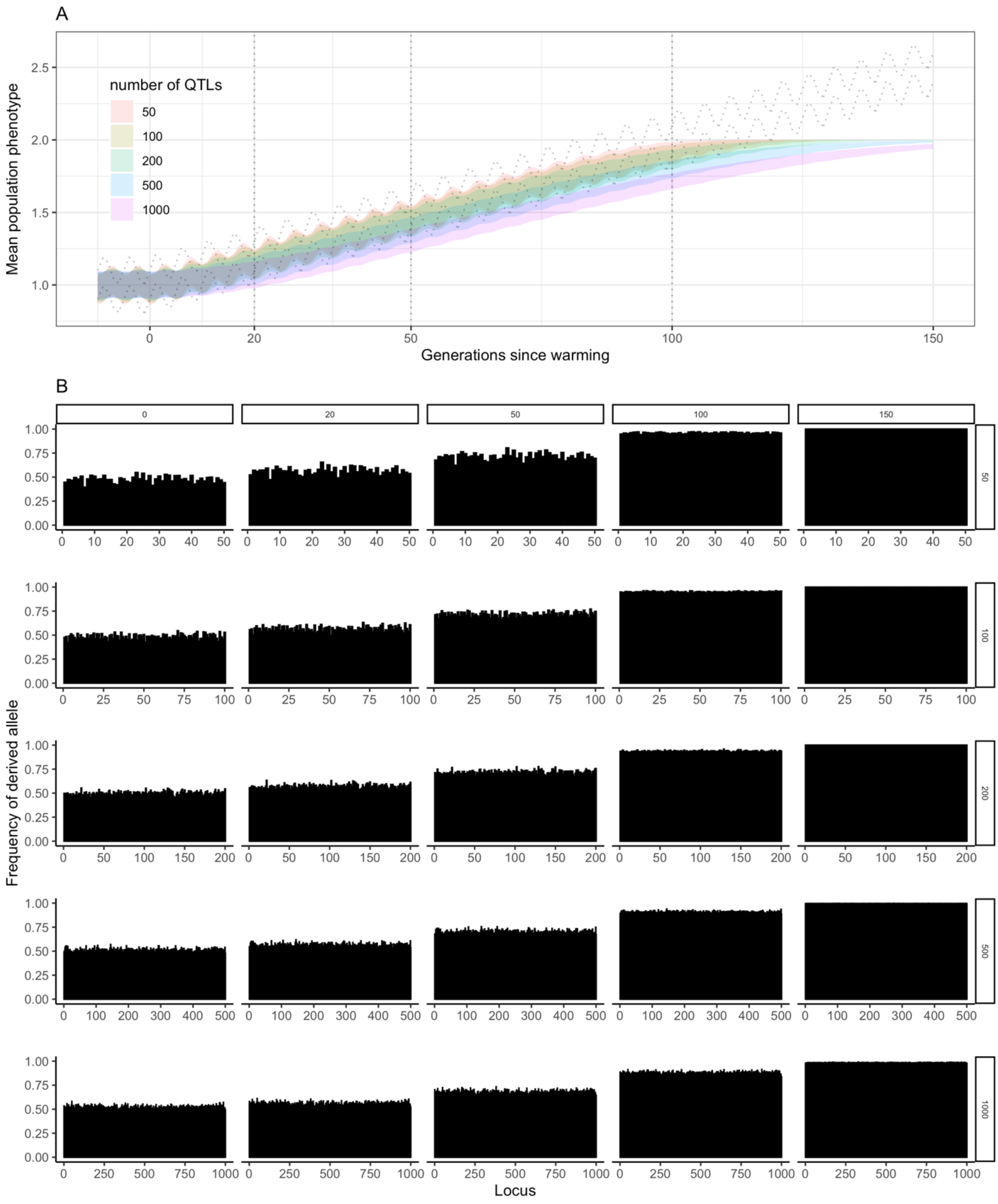


Supplementary figure 7. With increasing number of genes coding the trait under selection, adaptation is slower. The dashed line on panel A shows the range of environmental optima for the metapopulation, the coloured ribbons indicate the actual realised phenotypic range of the metapopulation for a trait coded by 50 to 1,000 loci. While a 50-QTL trait (in red) can respond readily to oscillations in the environment, as well as to progressive chronic changes (e.g. warming), a 1,000-QTL trait (in purple) is slower to respond (c.f. the waviness of the phenotype curve for the 50-QTL trait vs. the straightness of that of the 1,000-QTL trait). Accordingly, as warming progresses, the phenotypic change in the 1,000-QTL trait lags behind the environmental optimum and populations will be increasingly maladapted. This slow adaptation is unrelated to genetic diversity, and is simply a function of QTL number, as shown by the plots of the derived allele frequencies at various timepoints (Panel B). The allele frequency plots also illustrate the saturation of the adaptive capacity of the metapopulation until a point where all loci are fixed for the derived (beneficial) allele.

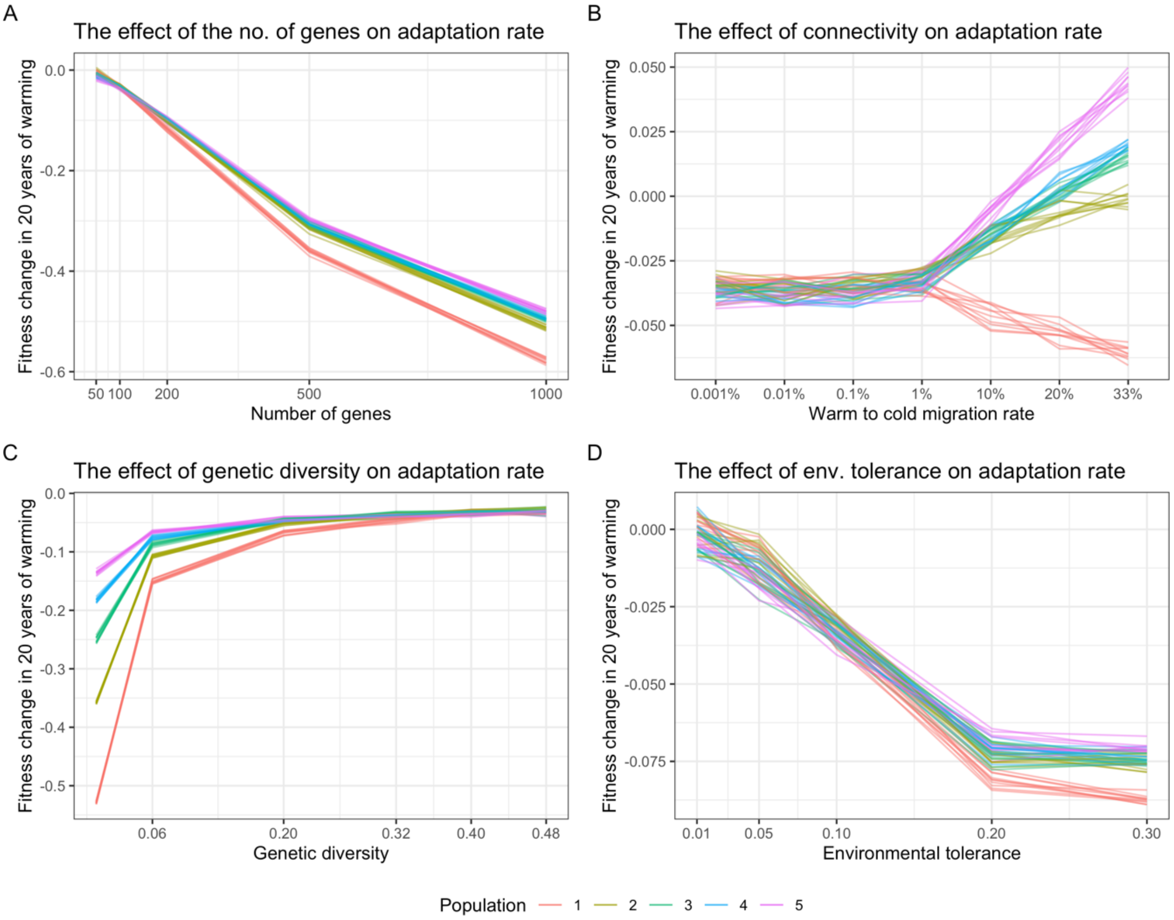


Supplementary figure 8. Fitness change in the first 20 years of warming in function of model parameters. A fitness change of 0 means that the population managed to keep pace with the rate of environmental change, while negative values show that the population became maladapted as the environment changed. Panel A: The higher the number of genes coding a trait, the slower adaptation is (see Supplementary Fig. 7), and therefore fitness decreases as the environment changes in 20 generations. Panel B: Higher rates of population connectivity increase the fitness for all but the warmest population, that is also the most upstream population in our stepping stone metapopulation model. The northernmost population suffers from outbreeding depression at increasing rates of connectivity; while all downstream populations benefit from the increased influx of warm-adapted genotypes. Panel C: At extreme low levels of genetic diversity (He < 0.06), adaptation is severely compromised. As diversity increases, adaptation rate follows a saturation curve. Panel D: Adaptation is slower with increasing levels of environmental tolerance , until a threshold value of 0.2 (10% of the maximum theoretical phenotypic trait value of 2.0).

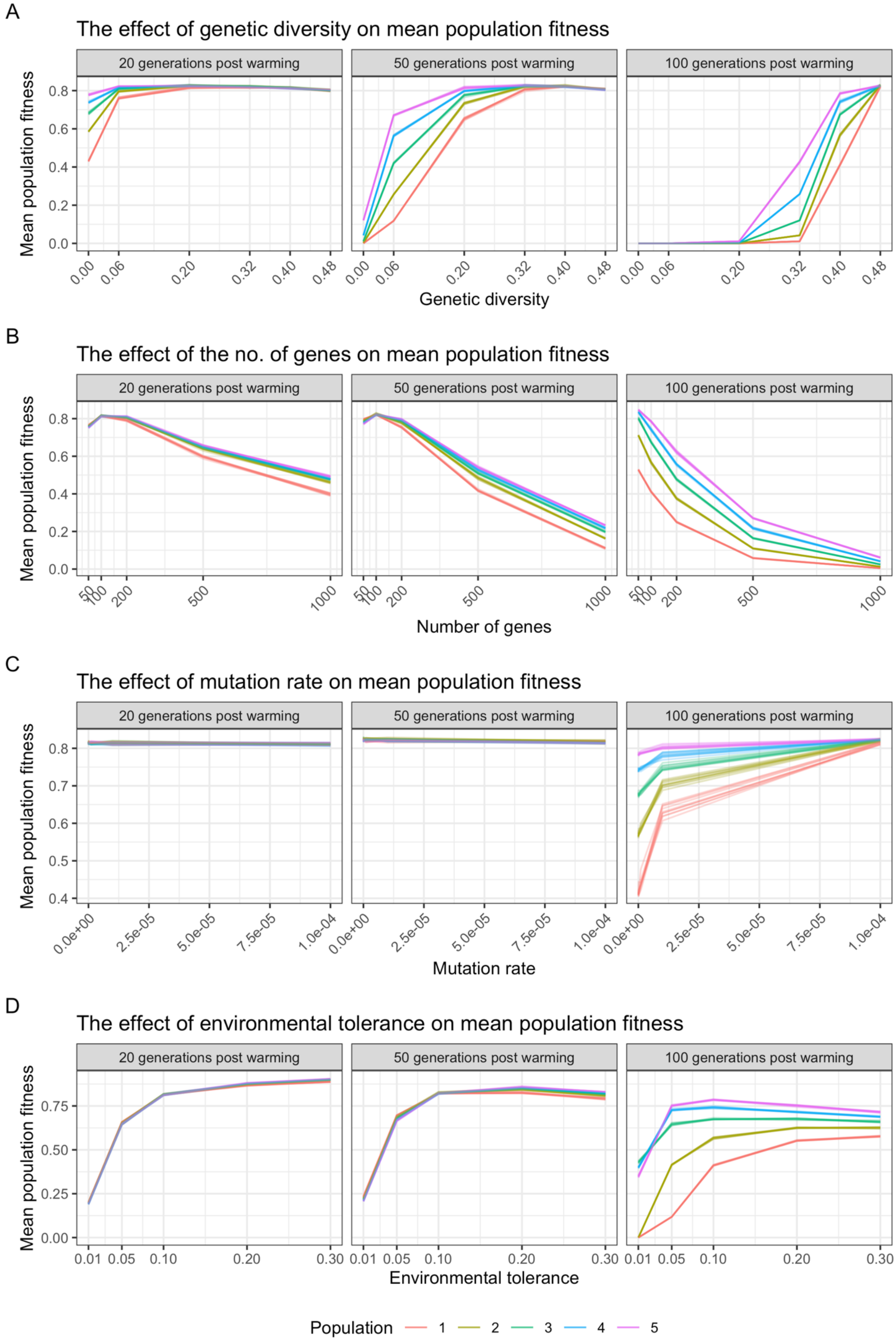
A
The effect of genetic diversity on mean population fitness
20 generations post warming
50 generations post warming
100 generations post warming
Mean population fitness
0.0
0.2
0.4
0.6
0.8
0.00
0.06
0.20
0.32
0.40
0.48
Genetic diversity
B
The effect of the no. of genes on mean population fitness
20 generations post warming
50 generations post warming
100 generations post warming
Mean population fitness
0.0
0.2
0.4
0.6
0.8
50
100
200
500
1000
Number of genes
C
The effect of mutation rate on mean population fitness
20 generations post warming
50 generations post warming
100 generations post warming
Mean population fitness
0.4
0.5
0.6
0.7
0.8
0.0e+00
2.5e-05
5.0e-05
7.5e-05
1.0e-04
Mutation rate
D
The effect of environmental tolerance on mean population fitness
20 generations post warming
50 generations post warming
100 generations post warming
Mean population fitness
0.00
0.25
0.50
0.75
0.01
0.05
0.10
0.20
0.30
Environmental tolerance
Population
1
2
3
4
5

Supplementary Figure 9. Fitness after 20, 50 and 100 generations of warming in function of model parameters. Panel A: Mean population fitness was higher at high genetic diversity, following a saturation curve characteristic to the population and the number of generations since warming. Panel B: Mean population fitness was generally lower as the number of loci coding a gene increased. Panel C: Mutation rate affected adaptation rates only 100 generations post warming, when the adaptive capacity of the metapopulation via standing genetic variation alone was depleted. Panel D: Mean population fitness was higher at high levels of environmental tolerance in the first 20 generations of warming. As warming progressed, 30 generations later the optimal environmental tolerance value was 0.2 for al populations. Close to the limits of adaptation, after 100 generations of warming populations 3-5 (on the colder end of the metapopulation) had a peak performance at 0.1 unit of environmental tolerance, whereas warmer populations (1 and 2) benefitted more from the highest environmental tolerance levels tested (0.3). There is an apparent contradiction between slower adaptation rate (Supplementary Fig. 8) and higher mean population fitness after 20 generations of warming when environmental tolerance was high. The explanation for this is that fitness values were already higher at the start of warming when environmental tolerance was high, compared to low environmental tolerance scenarios. Even with slower adaptation high environmental tolerance simulations reached higher overall fitness.

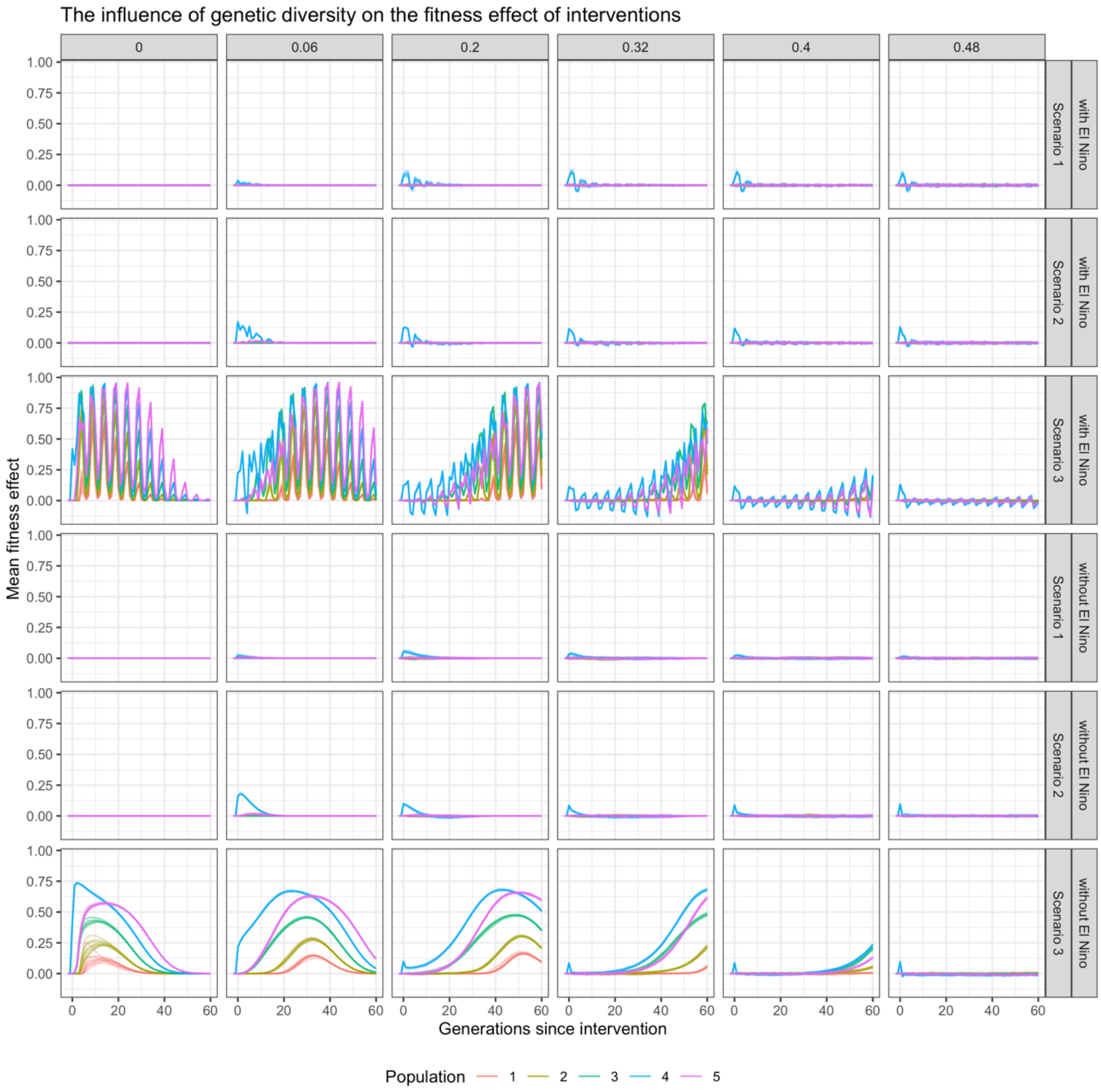


Supplementary figure 10. The influence of genetic diversity on the fitness effect of interventions, with and without ENSO-like environmental oscillation, for three different intervention scenarios (in facet rows). Genetic diversity (heterozygosity, He) values are in facet columns. The recipient of inoculum is population 4 (in blue), that is slightly colder than the mean of the metapopulation. At extreme low levels of diversity (He = 0), inoculum that carries only the standing genetic variation of the metapopulation (scenarios 1 and 2) yields no fitness effect because the metapopulation has exploited its adaptive capacity (evolutionary dead-end). At this low level of diversity, adding new genetic variation to population 4 (scenario 3) results in immediate and substantial fitness benefit that also spreads to other populations rapidly. The added variation expands the adaptive capacity of the metapopulation, and until all newly introduced alleles are fixed, there is a positive fitness effect compared to no intervention. Once the new alleles are fixed, the metapopulation's fitness plummets rapidly as it can no longer adapt to further environmental change, and its fitness becomes zero, similar to the no-intervention scenario. Above a heterozygosity value of 0.2, crossing the extremes populations (scenario 1) or artificially reshuffling the standing genetic variation (scenario 2) will create some fitness effect in the target population, but only temporarily. These 'fitness-ripples' fade away well within the first 20 generations from the deployment of intervention; and are not positive in all generations, depending on the ENSO-like phase. At the same time, adding new variation to increasingly diverse metapopulations results in smaller and smaller fitness effects, and similarly to other intervention scenarios, the effect is not always positive. At extreme high diversity (He > 0.48) the immediate effect of the introduction of new alleles to the metapopulation is indistinguishable from the effect of the other two intervention scenarios that work only with the standing genetic variation.

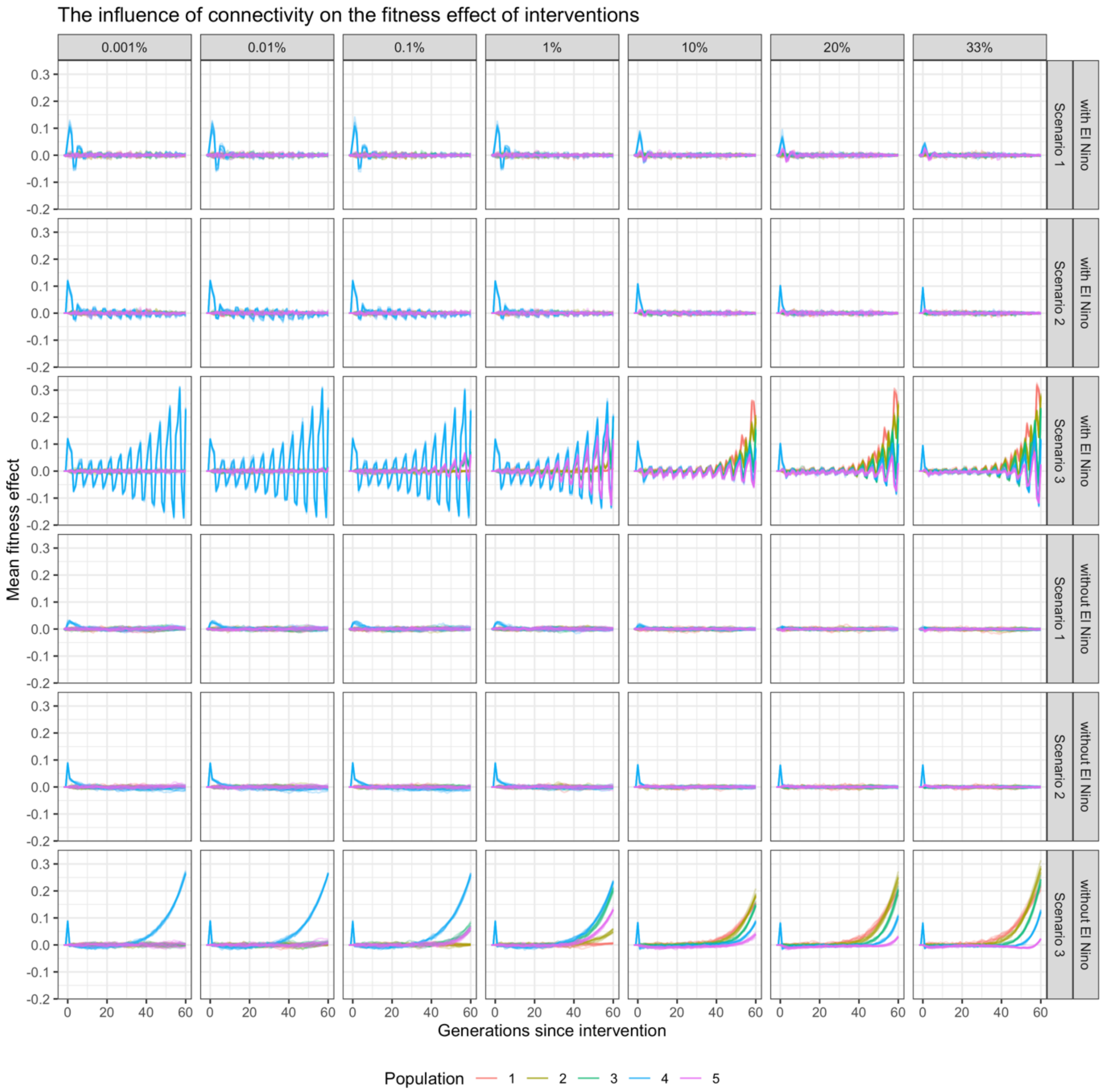


Supplementary Figure 11. The influence of connectivity on the fitness effect of interventions, with and without ENSO-like environmental oscillation, for three different intervention scenarios (in facet rows; Scenario 1 = crossing warm and cold populations; scenario 2 = re-shuffling the standing genetic variation within the metapopulation; scenario 3 = introducing novel genetic variation to the metapopulation). Connectivity rates tested are in facet columns. The recipient population is population 4 (in blue). Increasing rates of connectivity decrease the immediate fitness effect of the interventions on the target population, because the generations after the intervention are diluted by immigrants from other populations. For the same reason, the fitness effect of the intervention on other non-target populations increases with connectivity. This is particularly evident when new genetic variants are introduced as part of the intervention (Scenario 3). The new alleles introduced remain in the background for tens of generations, and play an important role as environmental conditions approach the theoretically possible phenotypic maximum of the metapopulation. As such, at high connectivity rates (larger than 10%) this intervention is most influential in the warmest population (population 1, red line), after 40 generations following the intervention.

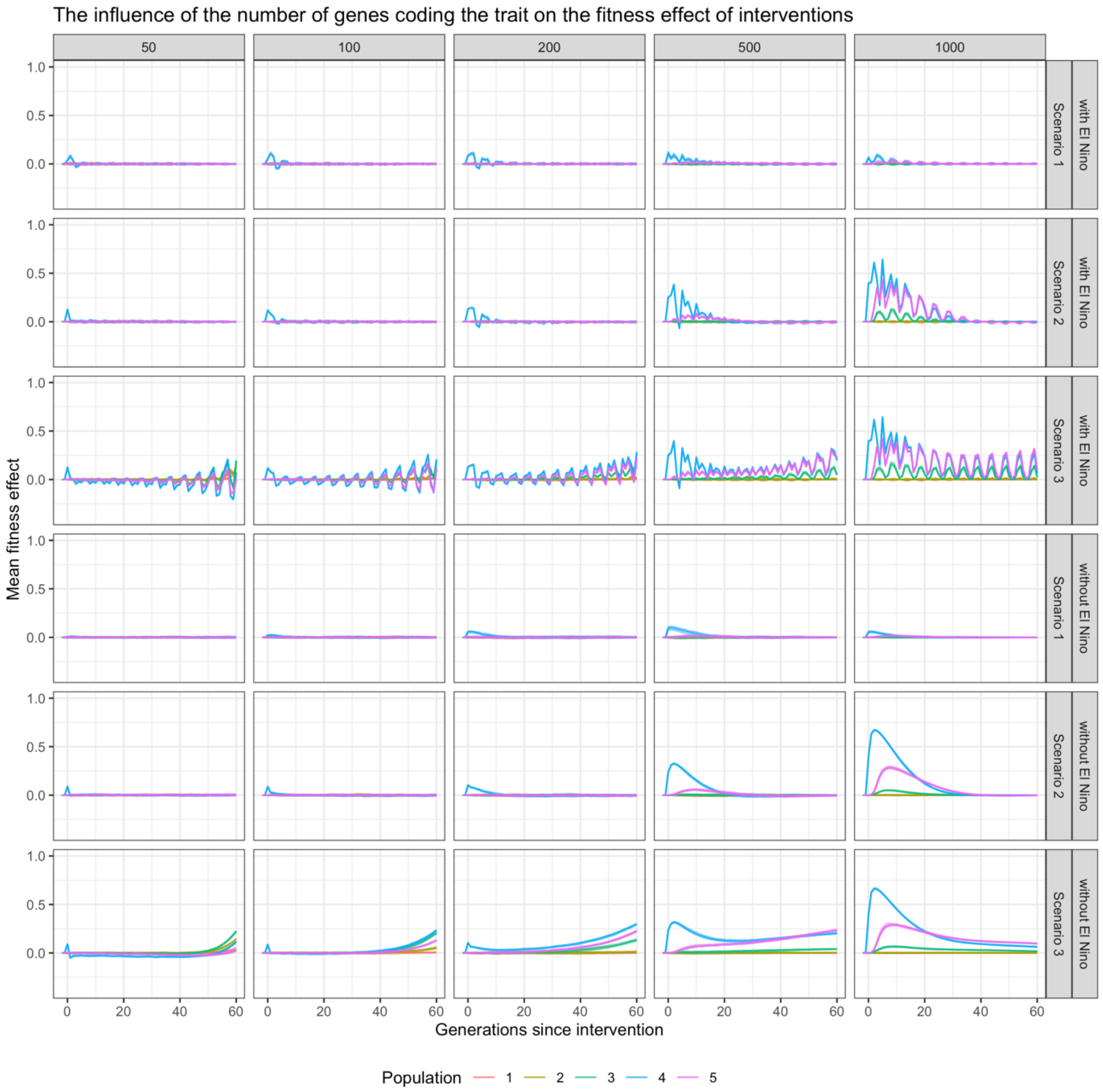


Supplementary figure 12. The influence of the number of genes coding the fitness-related trait on the fitness effect of interventions, with and without ENSO-like environmental oscillation, for three different intervention scenarios (in facet rows; Scenario 1 = crossing warm and cold populations; scenario 2 = re-shuffling the standing genetic variation within the metapopulation; scenario 3 = introducing novel genetic variation to the metapopulation). Numbers of loci tested in the simulations are in facet columns. The recipient population is population 4 (in blue). Interventions had a greater fitness effect when the trait under selection was coded by large numbers of genes. This is because the rate of natural adaptation is lower for highly polygenic traits (Supplementary Fig. 8), and therefore populations with highly polygenic traits were chronically maladapted at the time of intervention, 50 generations post warming. This also means that interventions had predominantly positive effect in simulations with highly polygenic traits, while they were either mainly negative, or partially negative for traits coded by lower numbers of genes.

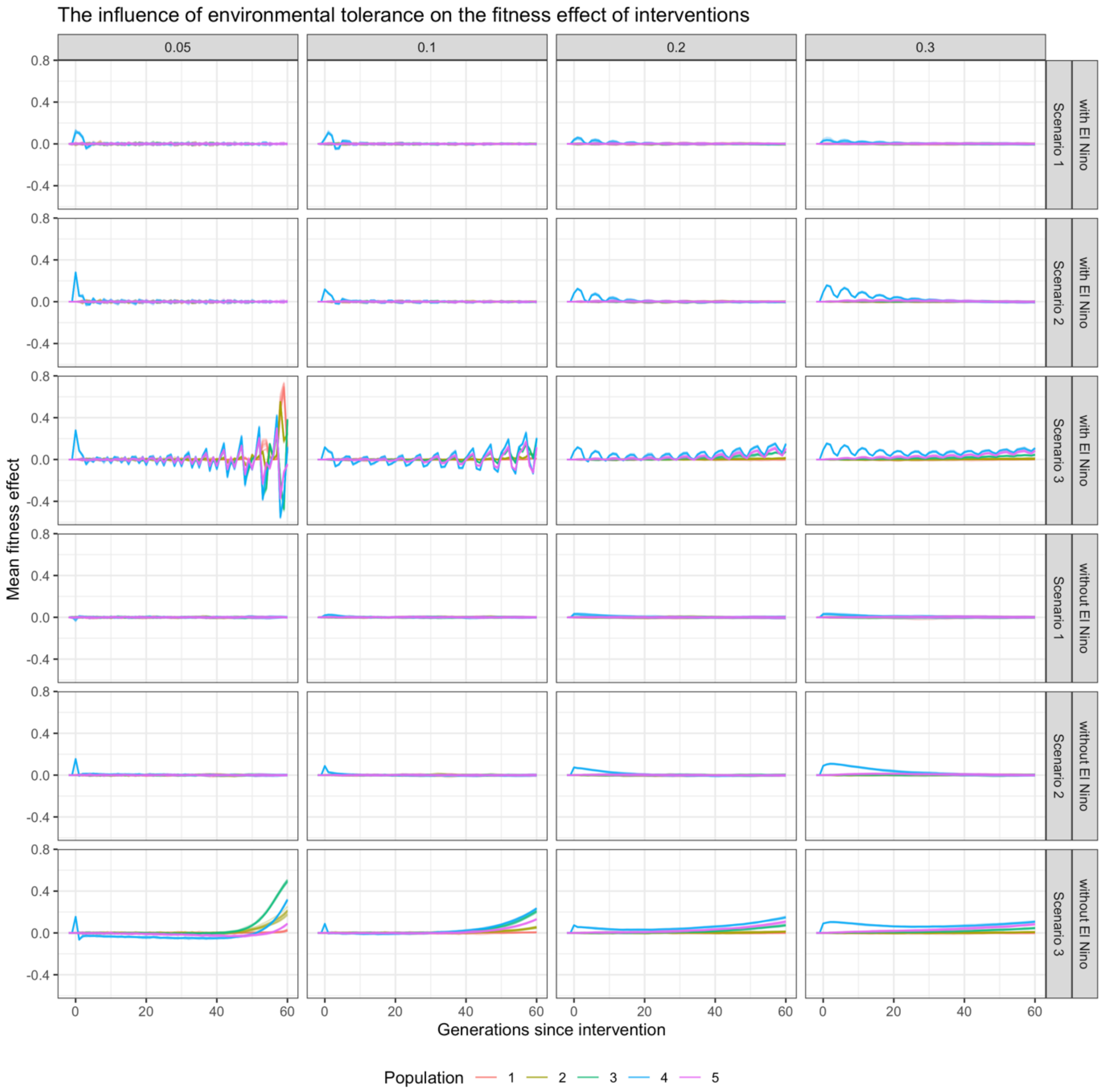


Supplementary figure 13. The influence of environmental tolerance (width /standard deviation/ of the fitness function) on the fitness effect of interventions, with and without ENSO-like environmental oscillation, for three different intervention scenarios (in facet rows; Scenario 1 = crossing warm and cold populations; scenario 2 = re-shuffling the standing genetic variation within the metapopulation; scenario 3 = introducing novel genetic variation to the metapopulation). Levels of environmental tolerance tested in the simulations are in facet columns. The recipient population is population 4 (in blue). The higher the environmental tolerance, the more positive and longer lasting the fitness effect of the interventions are.

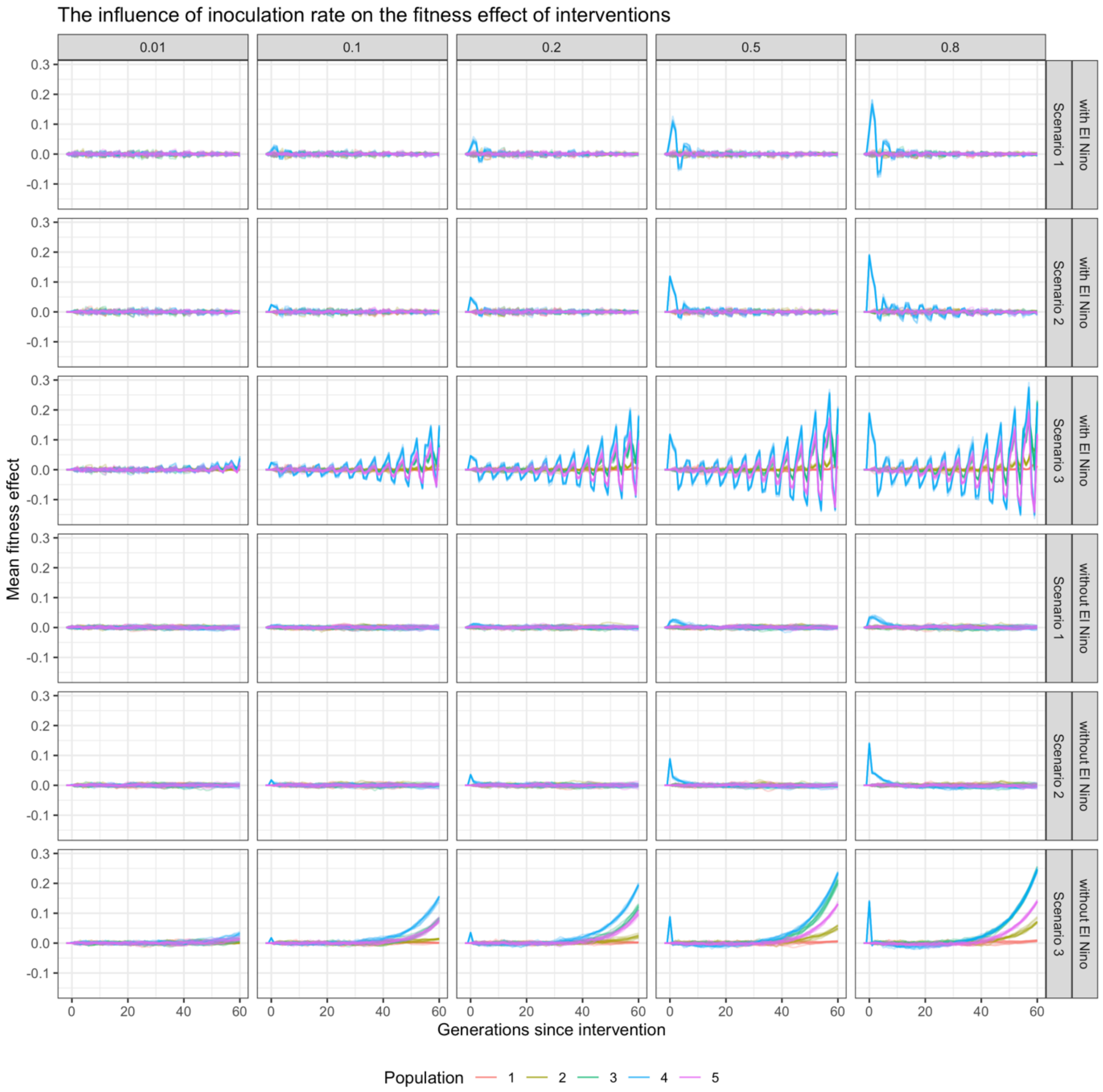


Supplementary figure 14. The influence of inoculation ratio on the fitness effect of interventions, with and without ENSO-like environmental oscillation, for three different intervention scenarios (in facet rows; Scenario 1 = crossing warm and cold populations; scenario 2 = re-shuffling the standing genetic variation within the metapopulation; scenario 3 = introducing novel genetic variation to the metapopulation). Inoculation ratios tested in the simulations are in facet columns. The recipient population is population 4 (in blue). The higher the inoculation ratio, the higher the effect of the intervention.
